# Continuity, not accuracy: how canopy approximations shape competition and selection in size-structured vegetation models

**DOI:** 10.64898/2026.11.18.767510

**Authors:** Lily P Dun, Daniel S Falster

## Abstract

Light competition drives ecological and evolutionary processes in forests. Real canopies are too complex to simulate in full, so stand models approximate them. We assessed three canopy approximations, the Perfect Plasticity Approximation (PPA), Flat Top and Deep Crown, in two stages. First, we compared their instantaneous light interception with a high-resolution reference model (MAESPA), focusing on competition among individuals and on vulnerability to self-shading. Second, we embedded them in a dynamic vegetation model and traced effects on stand development, demographic equilibrium and invasion fitness, the basis of evolutionary prediction.

Deep Crown, which allows leaf area to overlap between individuals, reproduced MAESPA’s continuous response of light capture to height. The PPA, and the Flat Top in stands of uniform height, instead produced a stepwise response around the point where a focal tree becomes taller than its neighbours. In the dynamic model, the PPA’s discontinuities were tolerable for stand development but shifted demographic equilibrium by an order of magnitude. Every runnable representation produced a continuously stable strategy (CSS), but under the PPA it was reached through a stepped, erratic selection gradient and was not robustly located. All approximations overestimated light capture under self-shading, increasingly so at realistic leaf area.

Our results show that continuity of the shade each plant casts as a function of height, rather than average accuracy or assimilation detail, determines whether an approximation can support evolutionary inference. Approximations adequate for stand-level light interception can mislead for competition and selection among individuals.

**HIGHLIGHTS:**

- Deep Crown tracked MAESPA light capture; PPA and Flat Top were stepwise
- PPA’s stepped shade shifted equilibrium seed rain tenfold in a dynamic model
- Continuity of cast shade, not assimilation detail, governs fitness landscapes
- All three approximations overestimate light capture when self-shading is strong

## 1 INTRODUCTION

Light competition is a dominant force structuring ecological and evolutionary processes in forests across the globe (Huston and Smith 1987; Falster and Westoby 2003; Purves and Pacala 2008). When trees become overtopped and shaded by others, the growth of smaller individuals is suppressed, driving an ecological process known as self-thinning, whereby the number of trees per unit area decreases as average individual size increases over time (Westoby 1984; Salas-Eljatib and Weiskittel 2018). When multiple species are present, competition between individuals also drives successional trajectories, where individuals from fast-growing species are progressively replaced by individuals from slower-growing species (Falster et al. 2017; Maxwell et al. 2019). Ultimately, competition among individuals leads to differential reproductive success, thereby influencing long-term vegetation dynamics, forest structure and composition (Falster and Westoby 2003; Fisher et al. 2010; Falster et al. 2017). Accurately capturing the effects of competition among individuals is therefore essential for any model seeking to predict demography, or predict the effects that environmental change will have on vegetation composition.

The dynamics of modelling light competition seem deceptively simple – leaves higher in the canopy shade those below them (Fisher et al. 2018). However, efficiently capturing the effects of competition between individual trees is a non-trivial computational challenge (Pacala and Deutschman 1995). Within current dynamical patch-based vegetation models, a variety of approaches are applied with varying complexity in how the canopy is represented, ranging from high-resolution 3D models to simple approximations. In high-resolution 3D models each tree is assigned a spatially explicit position (Pearcy and Yang 1996; Duursma et al. 2012; Duursma and Medlyn 2012; Vezy et al. 2018). These models then trace the daily and seasonal path of the sun in the sky; with 3D ray-tracing used to identify proportions of each crown the rays of light are hitting (Wang and Jarvis 1990; Medlyn 2004). However, the accuracy of this process requires considerable computing power, making it impractical for most dynamical models.

To reduce computational effort and thereby allow for modelling on larger temporal and spatial scales, multiple canopy approximations for light interception in stands have been developed, with three of these used within many current patch-based vegetation models (Fig. 1). The first is the Perfect Plasticity Approximation (PPA) (Strigul et al. 2008), used for example in FATES, which assigns cohorts to discrete canopy layers but resolves light through each crown (Fisher et al. 2015; Koven et al. 2020). The PPA assumes that tree crowns “perfectly” fill canopy gaps through phototropism, by reorienting their crowns towards higher light (Fig. 1**a**). For simplicity, the canopy is considered to be closed when it is full of tree crowns, thus the crowns organise into distinct canopy layers. In this approximation, there is a substantial and discontinuous drop in radiation at successively lower layers in the canopy, but no variation in radiation received among individuals in the same crown layer (Fig. 1**d**). This simplification greatly reduces the computational power needed, and can even allow analytical approximations of the size distribution, in certain circumstances (Strigul et al. 2008). Second, a “Flat Top” approximation is widely used, from the individual-based gap models (Bugmann 2001) to cohort-based models such as ED (Moorcroft et al. 2001; see Fisher et al. 2018). This approximation also ignores the horizontal spatial component of the canopy and assumes no vertical overlap between individual crowns, with the total leaf area of each tree distributed as a thin layer on top of the individual crowns (Fig. 1**b**). Hence, the canopy is represented as a series of infinitely thin layers, one for the crown of each individual. This causes a steep stepped vertical light profile, implying that there is a stepwise and sudden increase in light when an individual overtakes another (Fig. 1**e**). The third canopy approximation is the “Deep Crown” (Yokozawa and Hara 1995; Falster et al. 2011), which is similar to the Flat Top except that the leaf area of individuals overlaps in the vertical dimension (Fig. 1**c**). This allows the light profile to be a continuous function instead of a stepwise one and thereby reduces the benefit that would accrue to an individual that is slightly taller than surrounding vegetation, as only some of its leaves will rise above the other crowns (Fig. 1**f**). We therefore expect Deep Crown to be more accurate, but also more computationally expensive. While all three algorithms reduce computational expense, as approximations they will not perfectly capture the full details of a high-resolution model or true forest (Pacala and Deutschman 1995; Fisher et al. 2018; Braghiere et al. 2019).

**Fig. 1.**
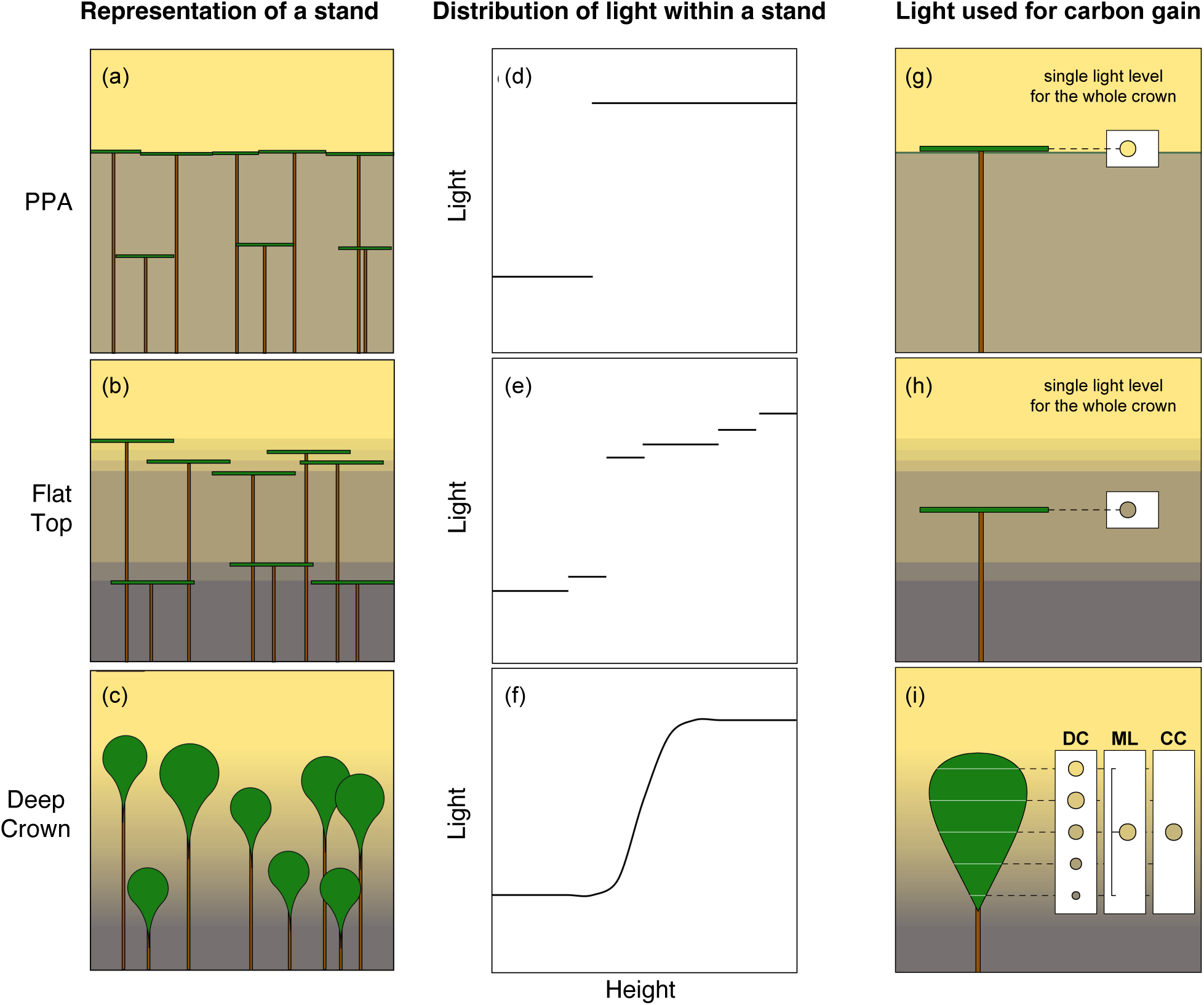
The two axes along which canopy approximations simplify light competition within a stand. Panels (**a**-**f**) show how a plant *casts* shade. Panels (**a**-**c**) show a hypothetical stand of trees under each approximation, which differ in how leaf area is distributed within crowns of individuals; yellow shading represents the different light levels resulting from the approximation. Panels (**d**-**f**) show the distribution of light with respect to height that occurs within a stand under alternate canopy approximations. Panels (**g**-**i**) show how a plant *reads* the light it sits in to compute its own carbon gain. The PPA and Flat Top assign a single light level to the whole crown (**g**, **h**). Panel (**i**) shows one crown casting the same continuous shade as in (**c**), read in the three ways compared in the dynamic experiments: the Deep Crown (DC) computes carbon gain at every layer and sums over crown depth, weighting by leaf area; Mean-Light (ML) averages light over crown depth, weighted by leaf area, and computes carbon gain once at that mean; Crown-Centre (CC) computes it once at the light at the crown centre. Circles show the light level(s) used, filled with the local light; circle area is proportional to the leaf area each represents.

Despite the potential for canopy approximations to alter competitive outcomes in vegetation models, the alternative approaches have not been quantified. Light interception and distribution have in general been evaluated in multiple studies (Stadt and Lieffers 2000; Beaudet et al. 2002; Song and Band 2004; Duursma and Mäkelä 2007; Roupsard et al. 2008; Kim et al. 2011; Fisher et al. 2018). However, Fisher et al. (2018) noted that how dynamic vegetation models partition light among competing canopies – including the canopy approximation and the details of light penetration and capture – controls the feedback between plant size and light acquisition, making it a pivotal component of these models. Braghiere et al. (2019) also showed that neglecting three-dimensional canopy structure, such as the clumping of foliage, leads an Earth system model to underestimate global photosynthesis. More recently, Eckes-Shephard et al. (2025) benchmarked nine vegetation demographic models against observations and found that, while the models captured the carbon content of mature forests, they diverged in post-disturbance regrowth, and shared compensating biases in growth and mortality. Yet none of these studies considers potential biases among individuals within stands. By comparison, alternative representations of leaf physiology have been extensively compared (Rogers et al. 2017). Understanding how well different models represent variation among individuals is essential, as this can fundamentally limit a model’s potential to inform the mechanics of species interactions, especially under environmental change. No study has yet traced how such individual-level biases propagate through a plant’s life cycle to alter the demographic and evolutionary predictions these models are built to make.

Here we ask how well three canopy approximations – commonly used in process-based vegetation models – capture individual competition: how much an individual tree intercepts light relative to the surrounding vegetation. We focus in particular on what happens as an individual moves from being shorter than the surrounding vegetation to being taller than it. Below, we use the term “point of transition” for the height at which a focal individual matches the surrounding stand. We hypothesise that the approximations will diverge substantially in their behaviour around this point, reflecting their differing complexity, with the more complex Deep Crown giving a more accurate representation of individual light capture.

We address this in two stages. First, we compare the *instantaneous* light interception of each approximation to a high-resolution 3D reference model, MAESPA (Duursma and Medlyn 2012), both for competition among individuals and for each approximation’s vulnerability to self-shading. MAESPA and its predecessors MAESTRO and MAESTRA have been widely applied and tested against observations of absorbed light, respiration, and carbon, water and energy fluxes across a range of species and systems (e.g. *Picea sitchensis*, *Betula pendula*, *Pinus taeda*, *Eucalyptus* plantations and *Coffea* agroforests) (Wang and Jarvis 1990; Wang et al. 1998; Luo et al. 2001; Medlyn 2004; Christina et al. 2016, 2017; Vezy et al. 2018; Braghiere et al. 2020), making it a reliable reference point. Because MAESPA is static, however, it cannot reveal how these instantaneous differences accumulate over a plant’s life. We therefore, in a second stage, embed the same kinds of canopy representation in a dynamic vegetation model and follow their consequences for stand development, demographic equilibrium, and the invasion fitness on which evolutionary prediction rests. In doing so we distinguish two axes along which an approximation can be simplified – how a plant casts shade on its neighbours, and how it reads the light it sits in for its own carbon gain – and ask which of these actually governs a model’s fidelity.

## 2 MATERIALS AND METHODS

We assessed three major canopy approximations (Fig. 1) in two stages. In the first, we compared the *instantaneous* light interception each approximation predicts to the output of a high-resolution model, MAESPA (Duursma and Medlyn 2012), establishing how faithfully each reproduces the light captured by an individual and how each responds to self-shading. Because MAESPA is a static model, it cannot show how these differences accumulate over a plant’s life. In the second stage we therefore embedded the same kinds of canopy representation in a dynamic vegetation model and followed their consequences for stand development, the demographic equilibrium of a population, and invasion fitness – the basis for evolutionary prediction (described under *2.7 Dynamic consequences* below).

### 2.1 Two axes of canopy approximation

Any canopy approximation makes simplifying assumptions along two distinct axes (Fig. 1). The first is how a plant *casts* shade: the vertical distribution of its leaf area, and hence the shape of the light profile it imposes on its neighbours (Fig. 1**a**-**f**). The second is how a plant *reads* the light it sits in to compute its own carbon gain (Fig. 1**g**-**i**). The two axes are independent – an approximation can coarsen one while resolving the other – and, as our results show, they are not equally consequential. The three approximations examined against MAESPA differ along both axes at once; we return to separating them in the dynamic experiments, where the distinction proves decisive. The two axes are also probed by different measurements. The shade-cast axis governs the light a plant intercepts and so is exposed by the MAESPA comparison of light interception below. The assimilation axis, by contrast, acts only through the nonlinear conversion of intercepted light to carbon gain (the saturating photosynthesis response): for a quantity linear in light, such as absorbed radiation, the leaf-area-weighted mean light and a full integration over crown depth coincide. A light-interception comparison is therefore structurally unable to separate the assimilation rules; that axis is taken up only in the dynamic experiments, where carbon gain is computed – which is why those experiments distinguish five representations where the MAESPA comparison resolves three.

### 2.2 MAESPA comparison of instantaneous light interception

For the first stage, each experiment involved simulating stand structure, then measuring light interception for focal trees within the same stand using four methods: the three approximations and the high-resolution control. We ran two experiments – competition among individuals, and vulnerability to self-shading within an individual’s crown (an overview is provided in Fig. **S1**). The same experiments were repeated under a variety of average stand conditions, as occurs in different vegetation types. In particular, we varied the density of trees to generate stands with different total shading, and the amount of variation in the heights of trees within the stand.

### 2.3 MAESPA description

As a control, we used the high-resolution model MAESPA (Duursma and Medlyn 2012). MAESPA simulates the fluxes of water, carbon and energy of individual trees within a stand at hourly time steps. Each tree is represented explicitly, with its own position and crown dimensions. Radiation is calculated at grid points within the crown of each target tree, separately for photosynthetically active radiation (PAR), near-infrared and thermal (long-wave) radiation (Duursma and Medlyn 2012). The radiation reaching each grid point depends on shading within and between crowns, given the sun’s position in the sky and the fractions of direct and diffuse radiation (Wang and Jarvis 1990). Direct, diffuse and scattered radiation are treated separately: scattering is approximated following Norman (1979), and at each grid point the leaf area is divided into sunlit and shaded fractions following Norman (1993).

### 2.4 Parameterisation of computational experiments

For the computational experiments, we simulated the spatial arrangement of trees in a stand with a given leaf area index (LAI, m² m^-2^) and height distribution. Each stand covered a ground area of 196 × 196 m, with trees placed on a regular square grid; the number of trees per row (26 to 89) set the stand density. A stand’s LAI is the leaf area of a tree divided by the ground area each tree occupies (the square of the tree spacing). To reduce edge effects (near sunrise and sunset), the two outermost rows of trees on each side formed a wall with high leaf area, short trunk height, high crown height and wide crown radius (Fig. **S2**). To further minimise edge effects, the canopy approximations used only trees more than 28 m from the edge of the plot (a central region of about 140 × 140 m). For experiments on individual competition and effects of self-shading, simulations were run for entire stands, with a focal tree located at the centre of each stand (Fig. 2**a,b**). Most parameters of the trees in the stand (not including the focal tree) were set as constant to reduce their effect on the outcome (Table **S1**). Each tree had a 15 m trunk, a crown 5.09 m wide and 6.36 m deep, and an individual leaf area (LA) of 27 m². This leaf area and crown depth lie within the range of *Eucalyptus* of similar height in the BAAD database (medians 32 m² and 6.2 m for trees 20–23 m tall), while the crown is wider than is typical [Falster et al. (2015); Fig. **S3**]. In the densest stands (tree spacing 2.2–2.5 m), neighbouring crowns overlap; MAESPA treats overlapping crowns as regions of higher leaf area density. Stands contained up to 7,921 trees, so we compiled MAESPA with its maximum number of trees raised from 1,500 to 10,000; with the default limit, MAESPA drops trees beyond the 1,500th and replaces the specified tree positions with its own grid. Each stand was driven by the same hourly meteorological data: an average day computed from two years of data (2003–2005) from the Tumbarumba research site (-35.66, 148.15) (Woodgate 2013). Fig. 2**f** shows the hourly light absorbed by an example focal tree in MAESPA, the same height as a sparse surrounding stand, across this average day.

**Fig. 2.**
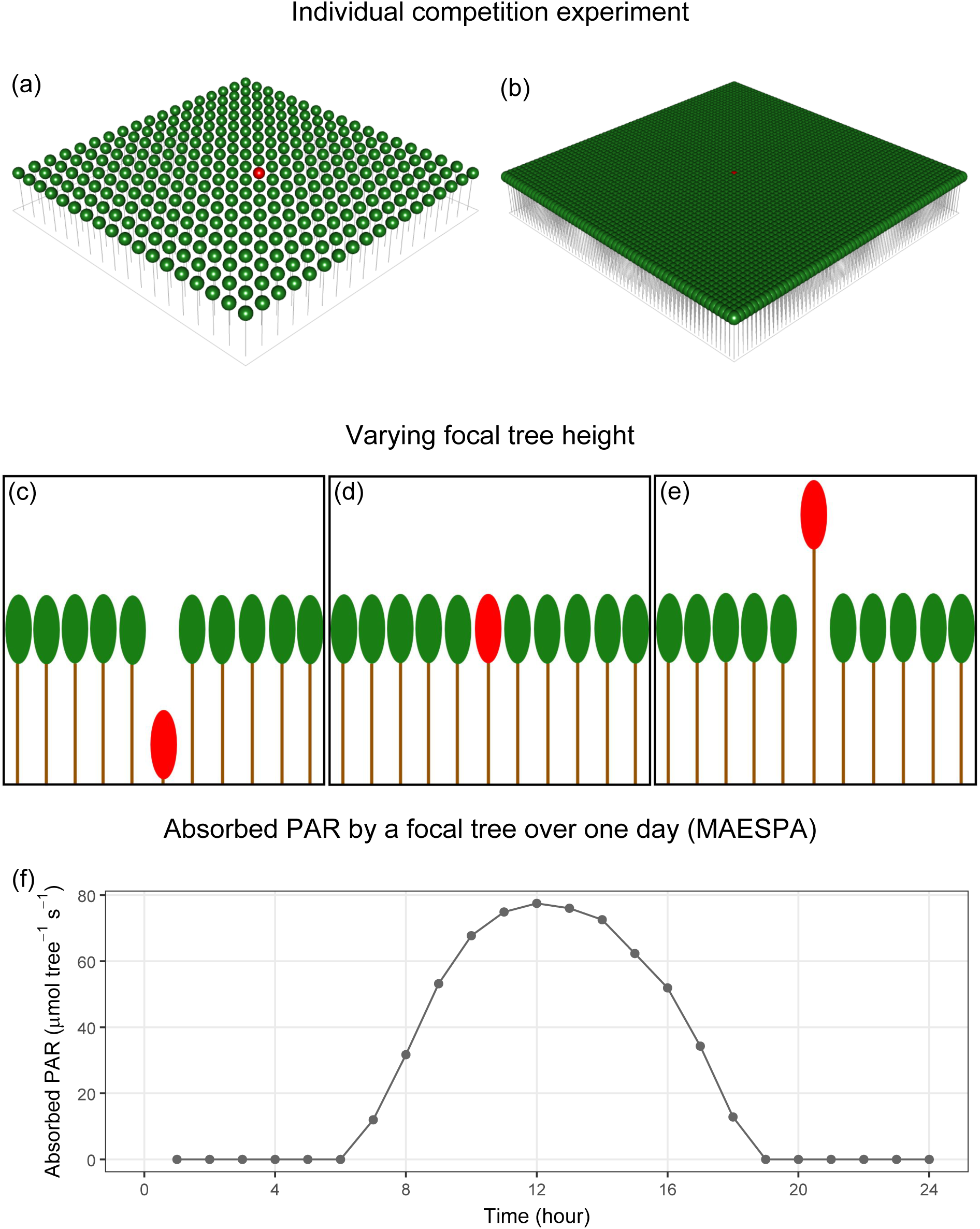
Examples of how the stands are represented in MAESPA. Panels (**a**, **b**) show the central region (about 140 × 140 m) of stands used in the individual competition experiment, with no variation in tree height and an LAI of 0.44 m² m^-2^ (**a**) and 5.44 m² m^-2^ (**b**); the focal tree is shown in red, and the edge-effect wall (Fig. **S2**) is not shown. Panels (**c**-**e**) represent different ratios of focal tree height to the surrounding stand: 0.01 (**c**), 1 (**d**) and 1.99 (**e**). Panel (**f**) shows the hourly absorbed photosynthetically active radiation (PAR) of a focal tree in the high-resolution model (MAESPA), in a stand with an LAI of 0.44 m² m^-2^, a CV of 0, and a focal-tree height ratio of 1, so that it is the same height as the surrounding stand. Time runs from hour 1 to hour 24 of a single day. Incoming radiation is the average day calculated from two years of data (2003–2005) at the Tumbarumba research site (latitude -35.66, longitude 148.15) (Woodgate 2013).

In each experiment, we tracked the light interception of one or more focal trees. Absorbed Photosynthetically Active Radiation (PAR) was calculated at hourly time steps and summed over the day to calculate the daily absorbed PAR for each focal tree. For each parameter combination, we simulated three replicate stands, which vary in the heights generated (drawn from a normal distribution, see below). We then averaged the daily absorbed PAR for each parameter combination.

#### 2.4.1 Individual competition experiment

The first computational experiment was conducted to understand how well the canopy approximations capture individual competition, compared to the high-resolution model. A total of 380 parameter combinations was considered, with 3 replicates each, leading to 1140 simulated stands. The LA of the focal tree was set to 0.1 m² to remove any self-shading within the focal tree, and thereby isolate the competitive effect. Self-shading is instead examined in the self-shading experiment below. The focal tree’s height in comparison to the rest of the stand was manipulated to 19 different ratios of the average trunk height of the stand, which was 15 m (Fig. **S1a**; Fig. 2**c-e**). We simulated variation in trunk height of trees in the stand by specifying a coefficient of variation (CV) (Fig. **S1a**). We used four different CVs, with 3 replicates each. Within each stand, heights were sampled from a normal distribution, with standard deviation determined from its CV. We used five stand densities, with LAI of 0.44, 1.42, 2.97, 4.28 and 5.44 m² m^-2^, spanning open stands in which crowns cover a third of the ground to closed stands in which crowns overlap.

#### 2.4.2 Vulnerability to self-shading experiment

The second computational experiment probed how vulnerable each approximation is to self-shading – the shading of a plant’s own leaves by others higher in its crown – by treating it as a known hard case and measuring each approximation’s sensitivity to it. The stand LAI was set as a constant 2.97 m² m^-2^, and the variation in trunk height as 0.1 CV. The LA of the focal tree was then altered to capture how the canopy approximations respond to self-shading within the focal tree (0.1, 0.5, 1, 10, 20 and 40 m²; Fig. **S1b**).

### 2.5 Implementation of canopy approximations

To implement the three canopy approximations, we took the morphological and physical parameters from each individual tree and stand, and applied the assumptions of each approximation. We then simulated light interception for the stand under each approximation.

#### 2.5.1 Perfect Plasticity Approximation (PPA)

For the PPA, we arranged the trees within the stand in descending order of their height and accumulated their crown projection areas from the tallest tree to the shortest. Following Strigul et al. (2008), canopy layers are filled in this order: a layer is complete once the cumulative crown area of the trees in it equals the ground area of the stand. Each tree was assigned to the layer in which its crown begins, and received the light below the leaf area of all complete layers above it. All trees in a layer therefore receive the same light, which is the main assumption of the PPA; in stands whose crowns cover less than the ground area, every tree is in the top layer and in full light.

#### 2.5.2 Flat Top Crown

For the Flat Top canopy approximation, we arranged the trees within the stand in descending order of their height then calculated the cumulative sum of the leaf area from the tallest tree in the stand to the shortest. We then divided the cumulative sum by the ground area occupied by the trees, to get the LAI above each tree.

#### 2.5.3 Deep Crown

For the Deep Crown canopy approximation, the crown of each tree within each stand was split into 50 horizontal slices, with leaf area distributed among slices in proportion to the volume of an ellipsoidal crown. We calculated how much leaf area would be in each slice of each crown in each of the simulations and arranged the slices in descending order of their associated height. We then calculated the cumulative sum of the leaf area from the tallest slice in the stand to the shortest. Therefore, each slice was assigned a height within the stand and the amount of leaf area above it in the stand.

### 2.6 Light model

Once the canopy for each stand was approximated, we calculated the light absorbed by the focal tree (or by each slice of the focal tree’s crown) from the leaf area above it. All three approximations used the same light model, so that differences among them reflect only how each represents the canopy. A single-component Beer’s law, in which light declines exponentially with the leaf area above (Cox et al. 1999; Kükenbrink et al. 2021), could not reproduce MAESPA across the range of stand densities (Fig. **S4**): in MAESPA, light declines steeply beneath the first leaf area above a tree, but much more slowly in dense stands, consistent with scattered light (Norman 1979) and light passing between crowns reaching the lower canopy. We therefore used an empirical two-component model, in which a fraction *w* of the incoming light attenuates quickly with leaf area and the remainder slowly:

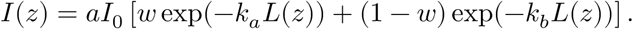

Here *I*(*z*) is PAR absorbed per unit leaf area at height *z*, *I*_0_ is PAR above the canopy, *L*(*z*) is the LAI above height *z*, *a* is the absorptance of leaves at the top of the canopy, and *k_a_* and *k_b_* are extinction coefficients for the two components (Table 1). We fitted these parameters to MAESPA using only focal trees at the two extremes of the competition experiment, in all 20 combinations of stand density and height variation: trees much taller than the stand (height ratio 1.99), which receive full light, and trees much shorter than it (height ratio 0.01), which sit beneath the whole stand. At these extremes all of the stand’s leaf area is either below or above the focal tree, so the Flat Top and Deep Crown give it the same leaf area above, and the fit cannot favour either approximation. We set *a* from the full-light trees and fitted *w*, *k_a_* and *k_b_* to the deep-shade trees by least squares, giving *a* = 0.75, *w* = 0.50, *k_a_* = 2.37 and *k_b_* = 0.42. Hourly PAR above the canopy, from the meteorological data, was converted from *μ*mol m^-2^ s^-1^ to MJ m^-2^ h^-1^ (Reis and Ribeiro 2020) and integrated over the day using trapezoidal integration. For the Deep Crown approximation, light interception was calculated for each of the 50 slices of the focal tree and weighted by the slice’s leaf area to obtain the absorbed PAR of the tree. The three stand replicates were then averaged. MAESPA outputs absorbed PAR per tree (MJ tree^-1^ d^-1^), so we divided these by the focal tree’s leaf area to express all estimates in MJ m^-2^ d^-1^.

**Table 1.**
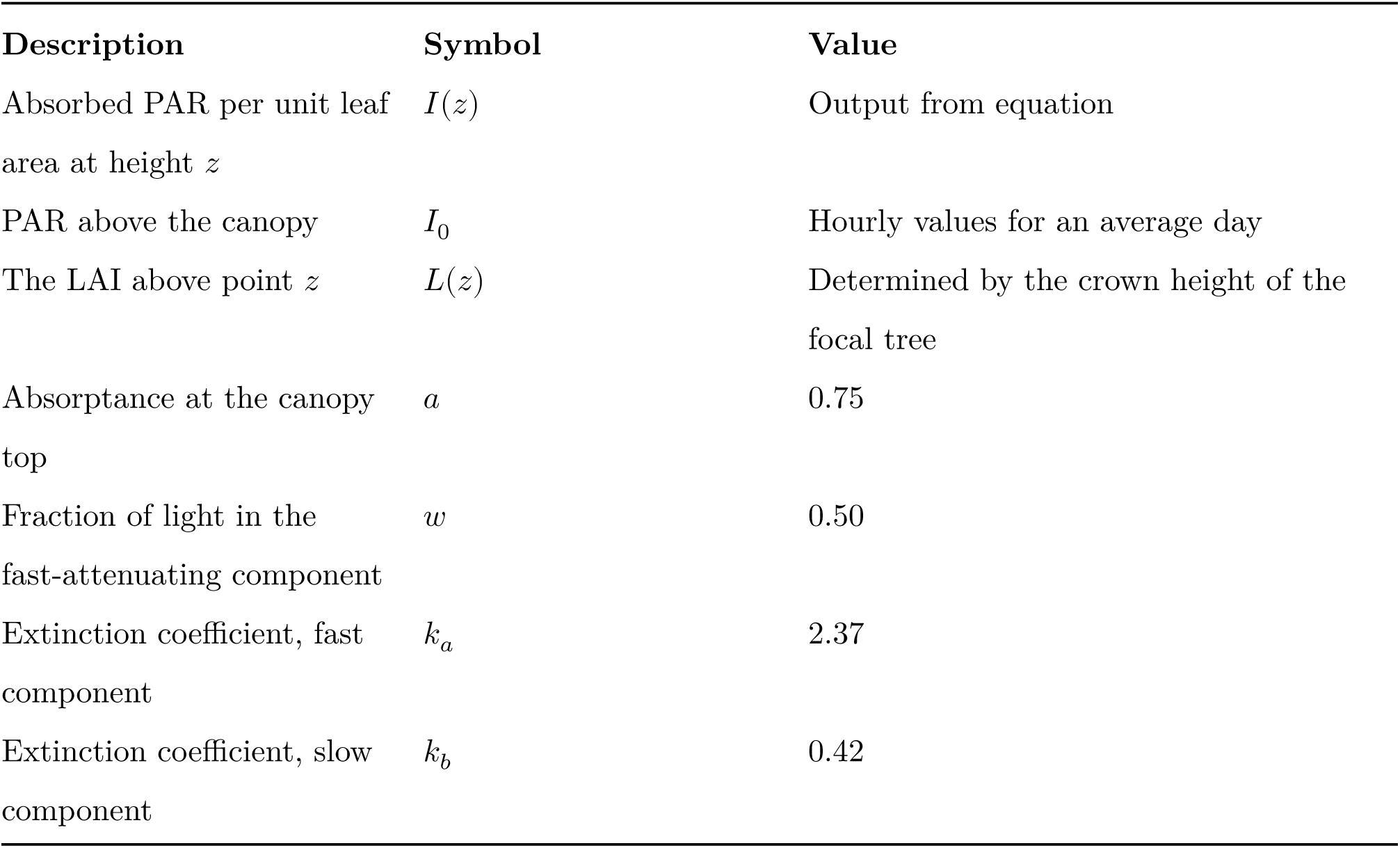
Symbols, descriptions, and values used in the light model. Parameter values were fitted to MAESPA (see Methods).

| **Description** | **Symbol** | **Value** |
|---|---|---|
| Absorbed PAR per unit leaf area at height *z* | *I*(*z*) | Output from equation |
| PAR above the canopy | *I*<sub>0</sub> | Hourly values for an average day |
| The LAI above point *z* | *L*(*z*) | Determined by the crown height of the focal tree |
| Absorptance at the canopy top | *a* | 0.75 |
| Fraction of light in the fast-attenuating component | *w* | 0.50 |
| Extinction coefficient, fast component | *k*<sub>*a*</sub> | 2.37 |
| Extinction coefficient, slow component | *k*<sub>*b*</sub> | 0.42 |

### 2.7 Dynamic consequences in a vegetation model

To trace how the instantaneous differences above accumulate over a plant’s life, we embedded the same kinds of canopy representation in plant, a trait-, size- and patch-structured dynamic vegetation model that resolves the growth, survival and reproduction of cohorts of plants competing for light within a patch (Falster et al. 2016). Each plant both contributes to and reads a shared patch light profile: its leaf area casts shade on the plants below it, and it reads the light at its own crown to compute carbon gain. The canopy approximations enter precisely as alternative rules for these two steps, which lets us separate the two axes of approximation. Deep Crown distributes leaf area continuously through the crown (following the profile of Yokozawa and Hara (1995) – a different crown shape from the ellipsoid used in the MAESPA comparison, but the same continuous, vertically overlapping distribution of leaf area) and integrates photosynthesis over crown depth; Mean-Light and Crown-Centre cast the same continuous shade but collapse the assimilation calculation to a single evaluation – of the leaf-area-weighted mean light, and of the light at the crown centre, respectively; the PPA reads a stepped, layered light profile – the field’s discrete canopy layers, whose boundaries we smooth (over 30% of each layer, the plant default) so the adaptive solver can integrate them (“PPA (smoothed)”), since the literal hard-step version used in large models (“PPA (hard step)”) does not run in continuous time; and the Flat Top collapses the shade a plant casts towards a thin layer at its crown centre, either as a hard step (“Flat Top (hard step)”) or smoothed into a continuous drop (“Flat Top (smoothed)”). A mapping between these names and the model’s settings is given in Table **S2**.

We simulated a single species (leaf mass per area, LMA = 0.0825 kg m^-2^), holding everything but the shading model fixed, and subjected each representation to three tests of increasing demand. First, *stand development*: we followed the leaf area index and size structure of a patch as it grew and selfthinned. Second, *demographic equilibrium*: we found the seed rain at which the population exactly replaces itself (net reproductive ratio R_0_ = 1), the self-consistent state a stand settles to. Third, *invasion fitness*: the lifetime reproductive output, log(R_0_), of a rare mutant growing through the light environment cast by an established resident (Metz et al. 1992; Falster et al. 2017), computed across a range of mutant LMA. Because invasion fitness depends on how the light a plant experiences changes as it grows – not merely the light at one size – it is the most demanding test, and a fitness landscape is useful for evolutionary analysis only if it is smooth. We computed each landscape under two seedings: all models at a common seed rain, isolating the effect of the shading model; and each model at its own equilibrium, so that each casts a self-consistent stand. From each landscape we located the singular strategy (the trait value at which the selection gradient vanishes) and classified it by testing both convergence stability – the sign of the selection-gradient slope through the singular point – and evolutionary stability – the sign of the fitness curvature at the singular point (Geritz et al. 1998). A singular strategy that is both convergence-stable and an uninvadable fitness maximum is a continuously stable strategy (CSS): a genuine evolutionary endpoint rather than a branching point. We took the steepness of the curvature and the smoothness of the gradient as measures of how robustly that endpoint is located.

plant integrates the patch with an adaptive, error-controlled solver, which requires the light a plant experiences to vary continuously with height. A literal discrete-layer profile is discontinuous and cannot be integrated; the PPA layers and the Flat Top box were therefore smoothed so that a stand could be solved. We also forced the hard-step PPA through a fixed-step integrator, to test whether it converges as the timestep is refined (Fig. **S8**).

### 2.8 Data analysis

The accuracy of the canopy approximations was evaluated by comparing the outputs to the high-resolution MAESPA outputs using root mean square error (RMSE) and R-squared analysis (R^2^) on each of the computational experiments. Because aggregate goodness-of-fit can mask the localised, qualitative differences that matter most for competition, we emphasise the shape of each approximation’s response alongside these summary statistics. Fit statistics use the 19 height ratios simulated in MAESPA. For the figures, the approximations were also evaluated at height ratios 0.005 apart, which requires no further MAESPA runs because each replicate stand is identical across focal heights; this shows where their steps occur, which the coarser sampling cannot resolve.

To create, run and analyse each of our experiments, we used R version 4.6.1 (R Core Team 2026), with the R package ‘Maeswrap’ (Duursma 2015) to set up the MAESPA simulations and read their outputs.

## 3 RESULTS

### 3.1 Individual competition

The competition experiment assessed how well the canopy approximations captured the effects of competition among individuals, by varying the height of a focal tree within a surrounding stand. The control from MAESPA shows that light interception of the focal tree decreases continuously in a logistic-like fashion, as the height of the focal tree decreases relative to the stand (Fig. 3). When it is much taller than other trees in the stand, light interception by the focal tree asymptotes at a maximum value, reflecting the maximum light available. When it is much smaller than other trees in the stand, light interception by the focal tree also asymptotes towards a minimum value, reflecting the accumulated shading above it. The rate of change in light interception by the focal tree is greatest around the point of transition. Supporting our hypothesis, we found that the Deep Crown approximation displayed a response similar in shape to MAESPA, whereas the PPA diverged, with a stepwise behaviour. In this stand, where tree heights vary (CV 0.25), the Flat Top tracked Deep Crown closely: its response steps up at the height of each neighbour, and with neighbours of varied height these steps are small. Deep Crown and Flat Top slightly underestimated light around the point of transition, where MAESPA’s response began to rise earlier and was less steep. The PPA instead overestimated light, both for shorter trees and, far more, for trees just taller than the stand, which it placed in full light (5.45 MJ m^-2^ d^-1^ at a height ratio of 1.05, against 2.25 in MAESPA).

**Fig. 3.**
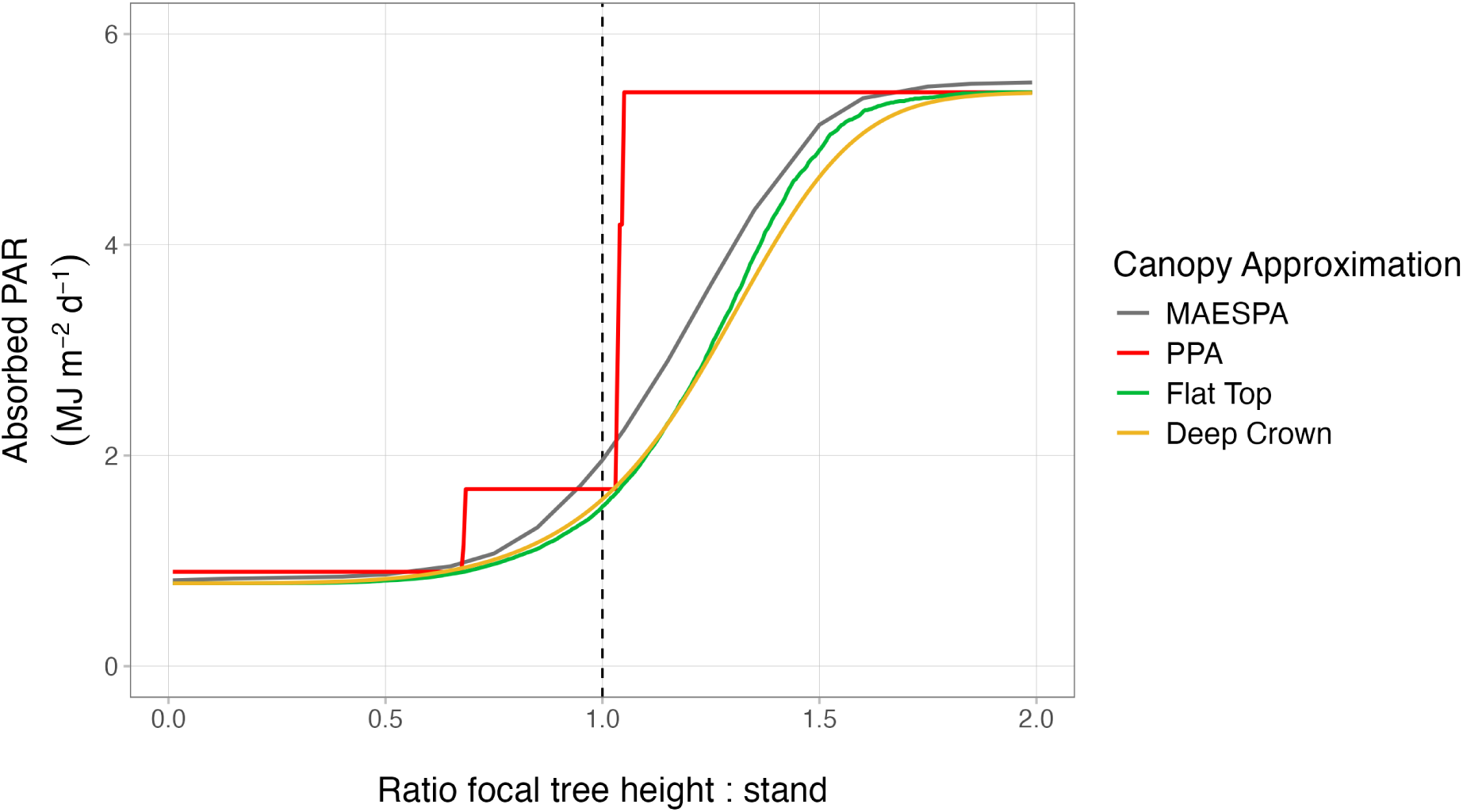
Results assessing the ability of three canopy approximations to capture the effects of individual competition. Lines show the light interception (absorbed PAR) for a focal tree for high-resolution model (MAESPA) and three different canopy approximations shown in Fig. 1. The x-axis is the ratio of the focal tree height to the stand mean, which is < 1 when the focal tree is shorter than the surrounding stand, 1 when the focal tree is the same height, and > 1 when the focal tree is taller than the average height of competitors (dashed line: the point of transition). MAESPA lines join the 19 simulated height ratios; the approximations are drawn at height ratios 0.005 apart, so that their steps appear where they occur. The simulated stand has a LAI of 2.97 m² m^-2^ and coefficient of variation of the height distribution is 0.25; it is the corresponding panel of Fig. 4.

Around the point of transition, the PPA diverged most strongly from the control (Fig. 3). Its response consisted of a few large steps: the focal tree received the same light across a wide range of heights, then jumped to full light once it reached the top canopy layer. The Flat Top showed a similarly sudden increase in stands where all trees had the same height (CV 0; Fig. 4): there, a marginally taller individual jumped from deep shade to full light (from 0.79 to 5.45 MJ m^-2^ d^-1^ between height ratios of 0.95 and 1.05, at LAI 2.97 m² m^-2^), whereas Deep Crown and MAESPA increased gradually (from 1.49 to 2.55 and from 1.80 to 2.90 MJ m^-2^ d^-1^ respectively).

**Fig. 4.**
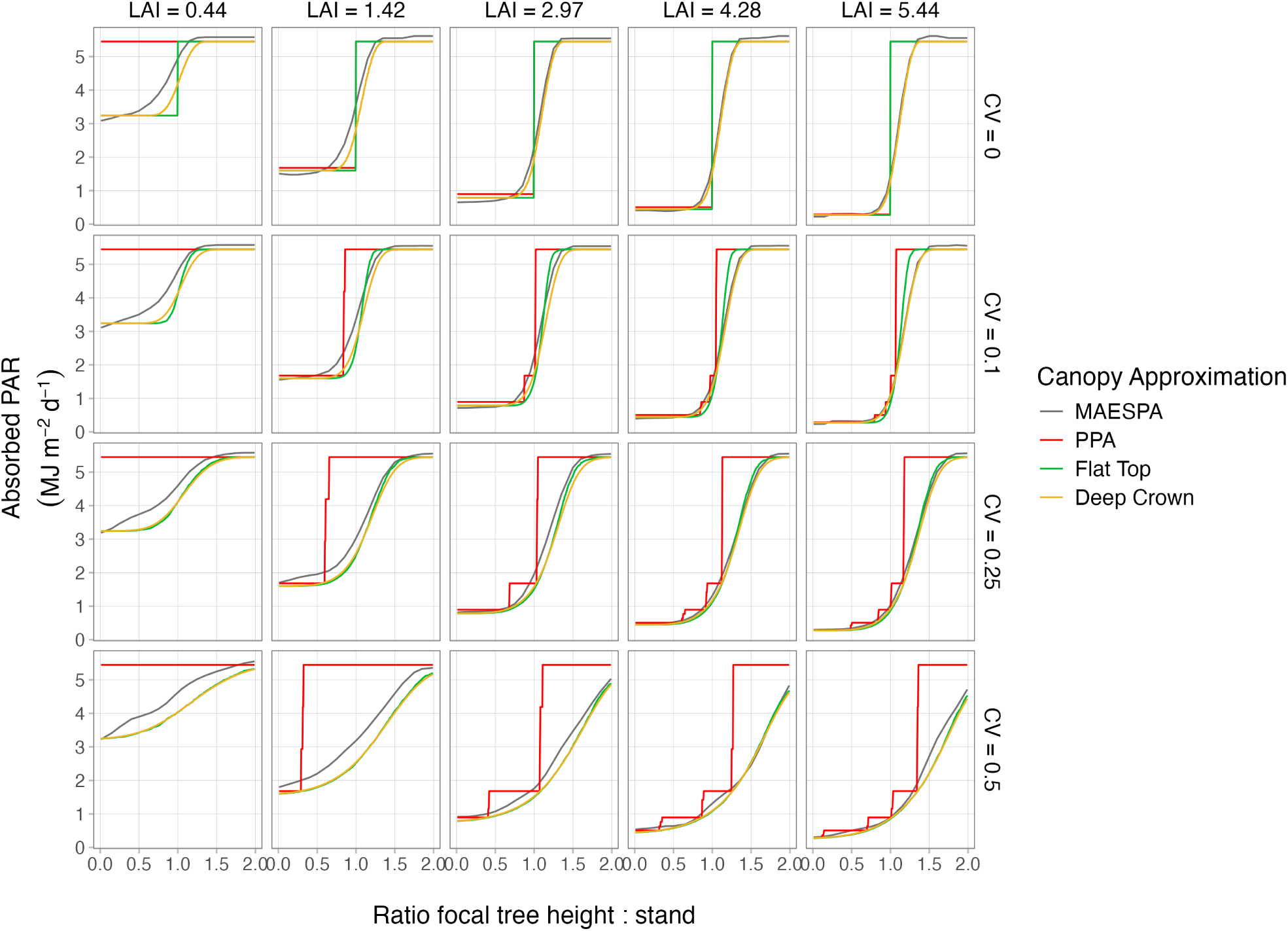
Results assessing the ability of three canopy approximations to capture the effects of individual competition under a variety of stand compositions. Lines show the light interception (absorbed PAR) for a focal tree for high-resolution model (MAESPA) and three different canopy approximations shown in Fig. 1. The x-axis is the ratio of the focal tree height to the stand mean, which is < 1 when the focal tree is shorter than the surrounding stand, 1 when the focal tree is the same height, and > 1 when the focal tree is taller than the average height of competitors. MAESPA lines join the 19 simulated height ratios; the approximations are drawn at height ratios 0.005 apart, so that their steps appear where they occur. Panels vary in the LAI and coefficient of variation of the height distribution within the stand (See methods for further details).

Over many scenarios of varying LAI and stand height variation, there was some behaviour that was consistent while there was variation (Fig. 4). The stepwise behaviour of the PPA was consistent through all scenarios but one: in the sparsest stands (LAI 0.44 m² m^-2^), where crowns cover only a third of the ground, the PPA’s canopy has a single layer, and every tree received full light regardless of its height. That of the Flat Top was pronounced in stands with little or no height variation (CV 0 and 0.1) but faded as height variation increased; at CV 0.25 and 0.5 the Flat Top was nearly indistinguishable from Deep Crown. In the denser stands (LAI 2.97–5.44 m² m^-2^), Deep Crown closely matched MAESPA (RMSE 0.13–0.27 MJ m^-2^ d^-1^ within each density). In the sparser stands (LAI 0.44–1.42 m² m^-2^), MAESPA’s light rose earlier and more gradually as the focal tree grew towards the height of its neighbours: trees well below the canopy received more light in MAESPA than in Deep Crown or Flat Top (e.g. 3.88 MJ m^-2^ d^-1^ in MAESPA against 3.29 for Deep Crown at a height ratio of 0.75, at LAI 0.44 and CV 0). Across all scenarios, Deep Crown fit MAESPA best (R^2^ = 0.99, RMSE = 0.29 MJ m^-2^ d^-1^), followed by Flat Top (R^2^ = 0.95, RMSE = 0.46) and PPA (R^2^ = 0.81, RMSE = 1.12). This whole-curve agreement nonetheless obscures the qualitative difference that matters for competition: only Deep Crown reproduces the continuous shape of the response in every scenario, whereas the PPA, and the Flat Top in stands of uniform height, misrepresent it precisely around the point of transition, where small height differences decide competitive outcomes.

### 3.2 Vulnerability to self-shading

In the individual competition experiment, the LA of the focal tree was set to a small value, to isolate competition among trees from self-shading within a tree. Here we instead set the focal tree’s LA to more realistic values, introducing a self-shading effect in MAESPA, and asked how vulnerable each approximation is to it. Because none of the approximations represent self-shading, all underestimated shading and overestimated light interception relative to MAESPA, and this divergence grew as focal leaf area increased (Fig. 5**a**; all six focal leaf areas are shown in Fig. **S5**). The vulnerability was shared by every approximation, including the best performer: when the focal individual has a LA of 0.1 m² so that self-shading is extremely minimal, Deep Crown fit MAESPA closely (R^2^ = 0.99, RMSE = 0.29 MJ m^-2^ d^-1^), as did Flat Top (R^2^ = 0.98, RMSE = 0.30 MJ m^-2^ d^-1^), followed by PPA (R^2^ = 0.90, RMSE = 0.75 MJ m^-2^ d^-1^). But at a realistic dense-crown LA of 40 m², Deep Crown retained a strong correlation yet was no longer accurate (R^2^ = 0.99, RMSE = 1.05 MJ m^-2^ d^-1^), particularly when the focal tree was taller than the stand, and the same held for Flat Top (R^2^ = 0.98, RMSE = 1.18 MJ m^-2^ d^-1^) and PPA (R^2^ = 0.90, RMSE = 1.56 MJ m^-2^ d^-1^). The divergence increased steadily with focal leaf area: for a focal tree much taller than the stand, the difference between Deep Crown and MAESPA grew from −0.09 MJ m^-2^ d^-1^ at 0.1 m² to 1.80 MJ m^-2^ d^-1^ at 40 m² (Fig. 5**b**). Because the approximations barely respond to the focal tree’s own leaf area, the same growth in error applies to all three.

**Fig. 5.**
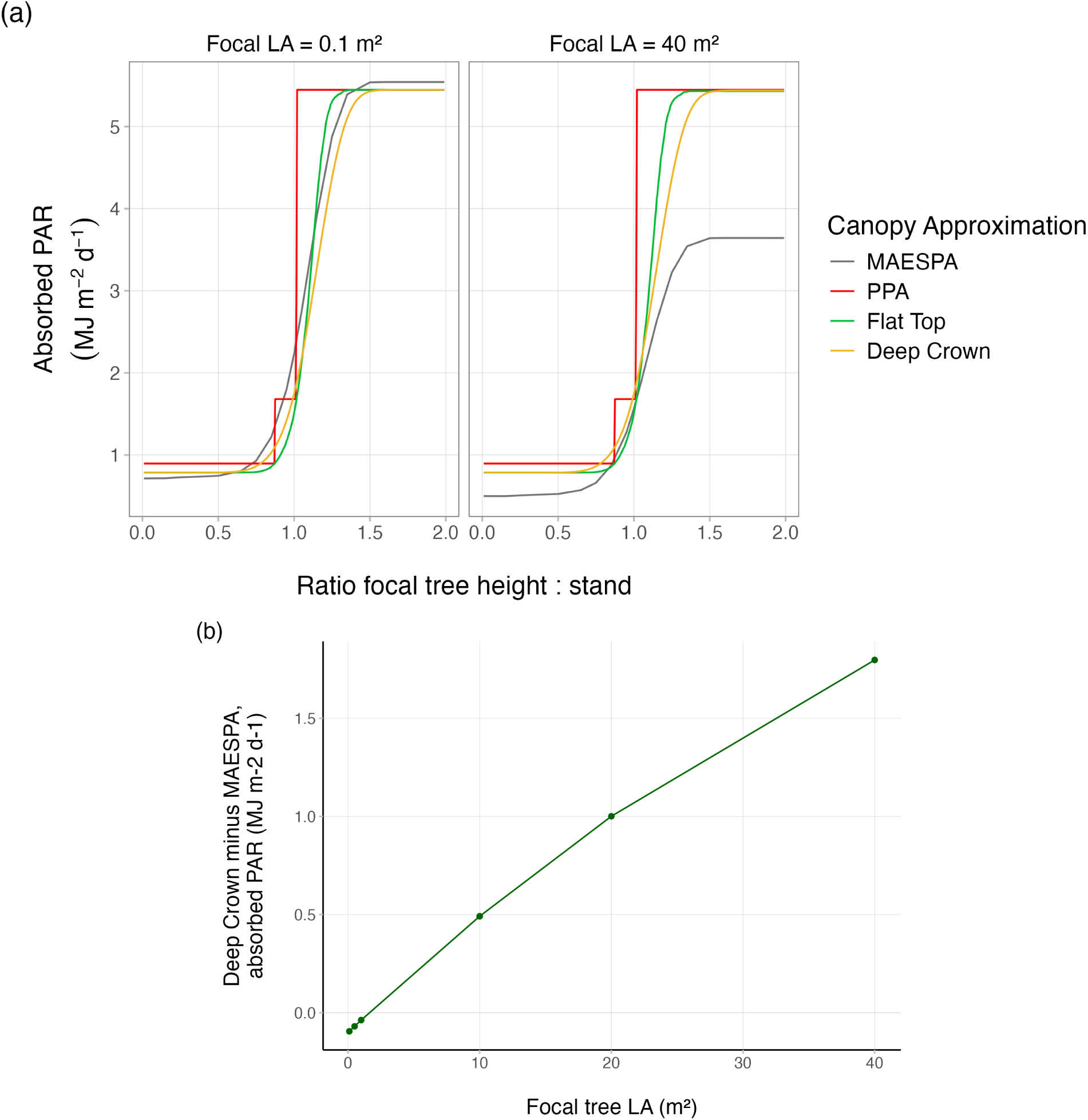
Vulnerability of the canopy approximations to self-shading. Panel (**a**): absorbed PAR for a focal tree against its height ratio to the stand, for MAESPA and the three approximations (Fig. 1), at a near-zero focal leaf area (0.1 m², where self-shading is negligible) and at a realistic dense-crown leaf area (40 m²). At 0.1 m² Deep Crown and Flat Top track MAESPA closely; at 40 m² all overestimate light capture because none represent self-shading, MAESPA’s curve falling well below them. Panel (**b**): the difference between Deep Crown – the most accurate approximation – and MAESPA, for a focal tree much taller than the stand (height ratio 1.99), as focal leaf area increases; even Deep Crown diverges progressively as self-shading grows. Both panels are from a stand with LAI 2.97 m² m^-2^ and CV 0.1. In panel (**a**), MAESPA lines join the 19 simulated height ratios and the approximations are drawn at height ratios 0.005 apart.

### 3.3 Dynamic consequences

#### 3.3.1 Stand development

Run forward in time, the four representations with a continuous light profile – Deep Crown, Mean-Light, Crown-Centre and the smoothed Flat Top – developed near-identical stands, with leaf area index rising and settling to a steady value (Fig. 6**a**). The PPA (smoothed) was the exception: its discrete light layers create sharp thresholds for cohort success, driving large, sustained oscillations in leaf area index as waves of recruitment and self-thinning pass through the stand. At this least demanding rung, then, the approximations are largely interchangeable in their size structure and vertical light profiles (Fig **S6**, Fig **S7**), save for the PPA’s layering artefact.

**Fig. 6.**
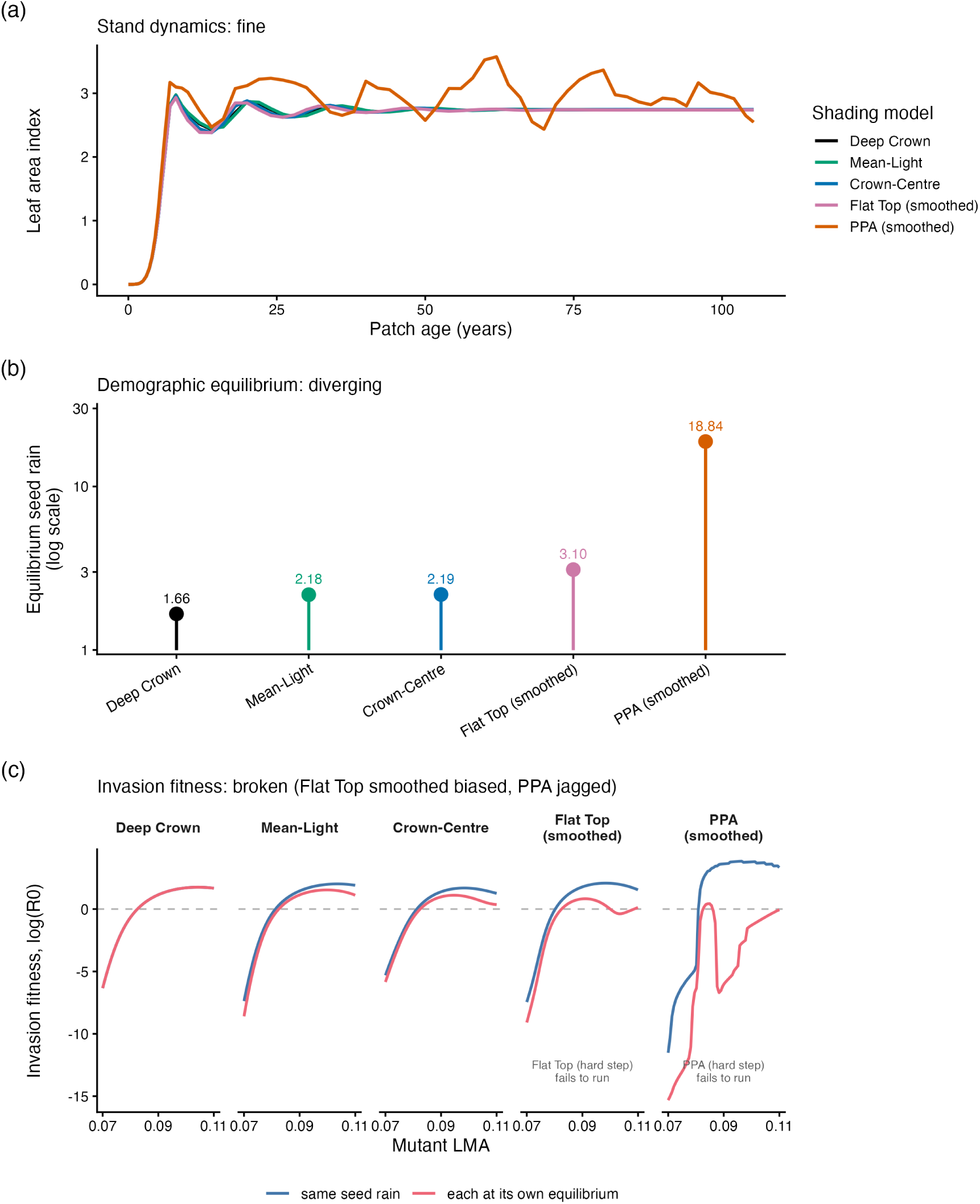
Consequences of the canopy representations in a dynamic vegetation model, shown as a ladder of increasingly demanding tests for a single species. (**a**) Stand development: leaf area index as a patch grows and self-thins; the continuous-competition models track closely while the PPA oscillates. (**b**) Demographic equilibrium: the seed rain at which the population replaces itself (log scale), spanning an order of magnitude across models. (**c**) Invasion fitness: the lifetime reproductive output, log(R_0_), of a rare mutant across mutant leaf mass per area, under two seedings (a common seed rain, and each model at its own equilibrium). The common seed rain is Deep Crown’s own equilibrium, so Deep Crown’s two curves coincide. Deep Crown, Mean-Light and Crown-Centre give smooth landscapes, the smoothed Flat Top’s landscape is distorted at its own equilibrium, and the PPA (smoothed) is jagged. Columns show the runnable models; the literal Flat Top (hard step) and PPA (hard step) make the light profile discontinuous and do not run. Model names follow the mapping in Table **S2**.

#### 3.3.2 Demographic equilibrium

The equilibrium seed rain – the rate at which a population exactly replaces itself – told a different story (Fig. 6**b**). Although the four models with a continuous light profile developed similar stands, their equilibria differed almost two-fold: 1.66 for Deep Crown, 2.18 (Mean-Light), 2.19 (Crown-Centre) and 3.10 for the smoothed Flat Top. The PPA (smoothed) was an outlier at 18.84 – more than ten times the Deep Crown value – so that across all five representations the equilibria spanned an order of magnitude. Because each model lets plants capture light slightly differently, each supports a different self-replacing density, and the PPA’s brighter layered light sustains a far denser, more productive stand. A difference this large, following from the shading assumption alone, shows that representations alike in their stand-development trajectories are not interchangeable once the population must sustain itself.

#### 3.3.3 Invasion fitness

Invasion fitness was the most discriminating test (Fig. 6**c**). Deep Crown, Mean-Light and Crown-Centre produced smooth fitness landscapes under both seedings, whereas the PPA (smoothed) was jagged – and at its own, much denser equilibrium, wildly so, with a non-monotonic dip where small trait changes flip which canopy layer a mutant ends in. The smoothed Flat Top ran, but at its own equilibrium its landscape was distorted, dipping below zero again at high LMA. The literal field-standard discretisations did not run at all: the Flat Top (hard step) and the PPA (hard step) each make the patch light profile discontinuous, which the adaptive solver cannot integrate.

Classifying the singular strategy in each landscape sharpened this picture (Fig. 7; Table **S3**). Every representation produced a singular strategy that was both convergence-stable – the selection gradient crossed zero with negative slope (Fig. 7**a**) – and an uninvadable fitness maximum, with negative fitness curvature: a continuously stable strategy (CSS), and so a genuine evolutionary endpoint rather than a branching point. The existence of an endpoint was thus qualitatively preserved across approximations; what differed was where it lay and how robustly it was reached. The endpoints disagreed by about 15% in trait value, the singular LMA falling monotonically from 0.101 (Deep Crown) through 0.098 (Mean-Light), 0.092 (Crown-Centre) and 0.090 (Flat Top, smoothed) to 0.086 (PPA, smoothed) (Fig. 7**b**) – similar fitness-landscape shapes settling on different optima because each representation reaches a different equilibrium density. The fitness curvature at the endpoint also steepened about twentyfold, from about −1.2 × 10^4^ (Deep Crown) to −2.5 × 10^5^ (PPA, smoothed). Combined with the PPA’s stepped, erratic selection gradient (Fig. 7**a**, inset; Fig. 6**c**), this means its CSS, though formally present, is not robustly located: because small trait differences flip a mutant between canopy layers, the selection gradient between nearby strategies – the quantity that determines which way a trait evolves – is unreliable even where an endpoint exists. Every other representation, including the smoothed Flat Top, placed the CSS through a smooth gradient and so located it robustly; the smoothed Flat Top’s endpoint (0.090) lay close to that of Crown-Centre (0.092). Among these smooth representations, the endpoints differ mainly through their different equilibrium densities, not through the continuity of the gradient.

**Fig. 7.**
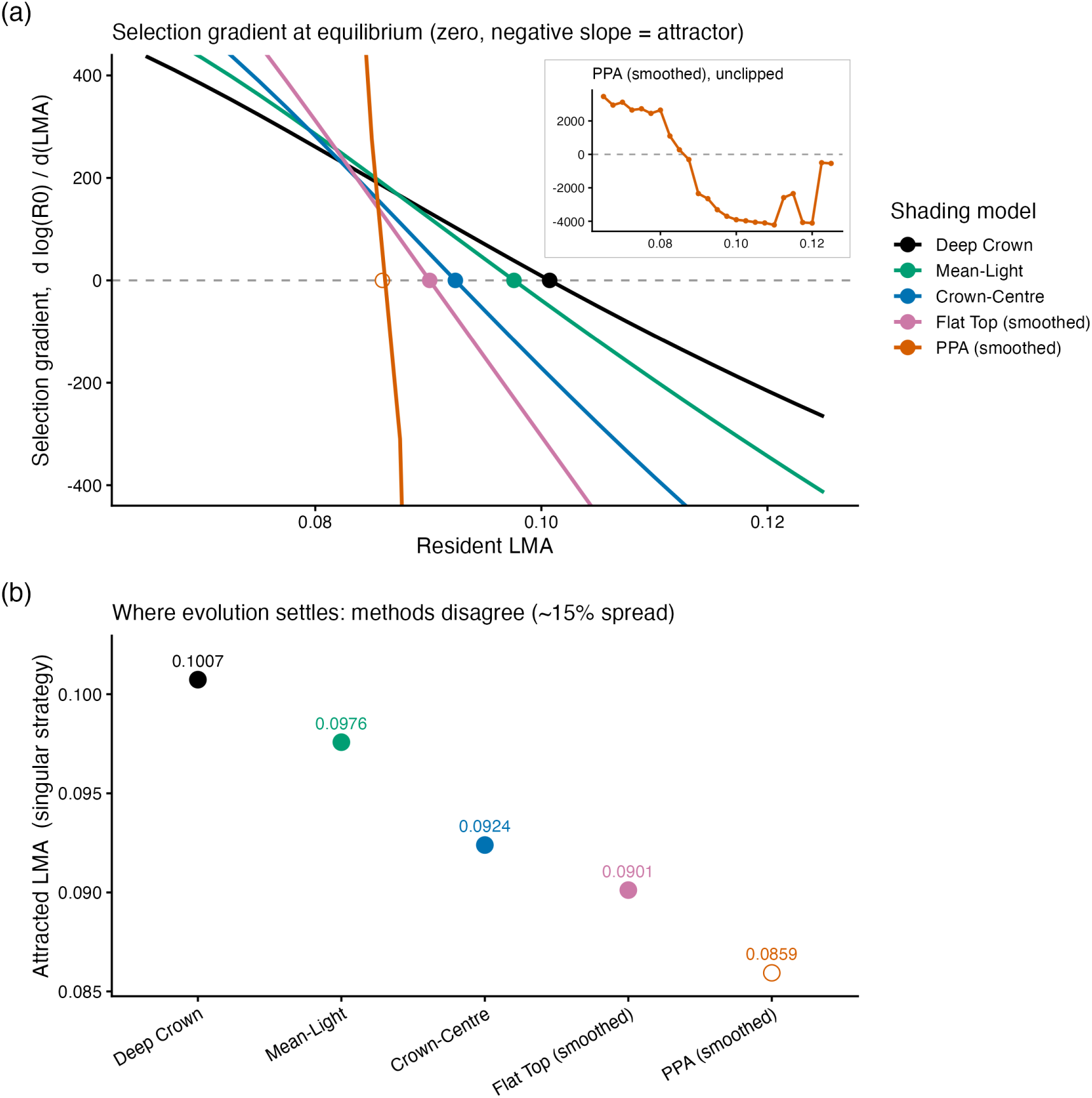
Evolutionary endpoints under each canopy representation. (**a**) The selection gradient (d log(R_0_) / d LMA) as a function of resident LMA; a zero-crossing with negative slope is a convergence-stable singular strategy. The main panel is clipped at ±400 so that the smooth crossings are legible; the inset shows the PPA (smoothed) gradient over its full range. (**b**) The resulting singular strategy – the trait value evolution settles on – for each representation. All five are continuously stable strategies (CSS): convergence-stable and uninvadable fitness maxima, hence genuine evolutionary endpoints. Yet they disagree by about 15% in trait value, because each representation reaches a different equilibrium stand density. The PPA’s selection gradient is stepped and erratic because of its discrete layers (inset; open symbol), so its endpoint, though a CSS, is not robustly located. Model names follow the mapping in Table **S2**.

That Mean-Light and Crown-Centre – which approximate a plant’s own assimilation as crudely as the Flat Top, by a single evaluation – nonetheless give smooth, usable fitness landscapes, which points to where the problem lies: the decisive difference is the shape of the shade a plant casts, not how it reads its own light. We make this explicit by comparing representations that share the crown-centre assimilation but differ in their competition profile (Fig. 8**a**). Crown-Centre casts a smooth shade that builds continuously down through the whole crown, whereas the Flat Top concentrates it: in a single step (hard step), or in a continuous drop compressed into the upper part of the crown (smoothed). At each model’s own equilibrium, the Crown-Centre landscape keeps the single broad maximum of the Deep Crown reference, although it peaks lower and at a lower LMA; the smoothed Flat Top landscape instead dips below zero again between LMA of about 0.10 and 0.11 before rising (Fig. 8**b**), a distortion absent from every representation that casts the smooth Yokozawa shade. The hard step gives no landscape at all, because its discontinuous shade leaves the stand unsolvable. Continuity of the cast shade therefore determines whether a model can be solved, and whether its fitness landscape keeps its shape; how a plant reads its own light shifts where the optimum lies, but not whether it is well defined.

**Fig. 8.**
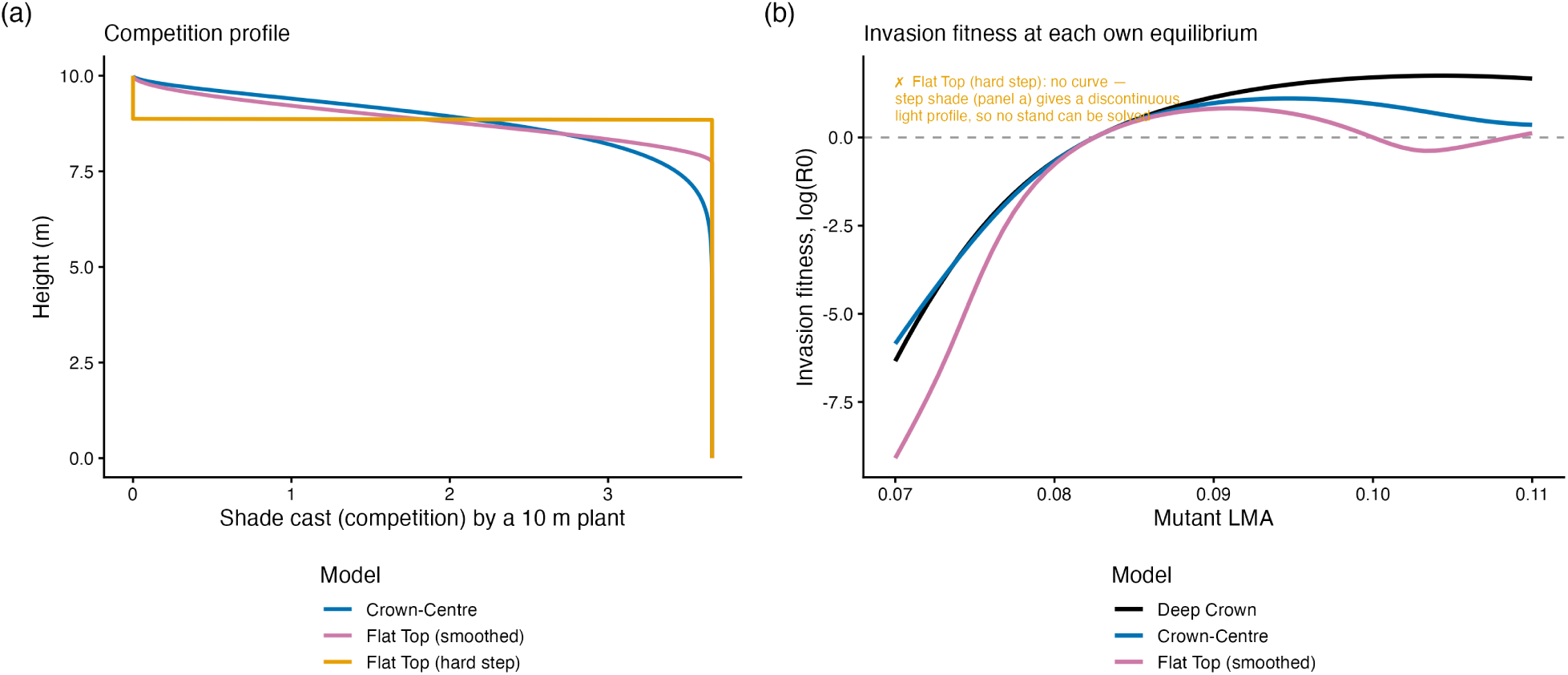
The shape of the cast shade, not the assimilation rule, governs fitness. (**a**) The shade (competition) cast as a function of height by a single 10 m plant under three representations that share the same crown-centre assimilation: Crown-Centre casts a smooth shade that builds continuously down through the crown, the Flat Top (hard step) concentrates it in a single step, and the Flat Top (smoothed) in a continuous drop compressed into the upper crown. (**b**) Invasion fitness landscapes, each model at its own equilibrium: the Crown-Centre landscape keeps the single broad maximum of the Deep Crown reference (though lower, and at lower LMA), whereas the smoothed Flat Top landscape dips below zero again at high LMA; the Flat Top (hard step) yields no landscape because its discontinuous shade leaves the stand unsolvable.

## 4 DISCUSSION

Our study examined three common canopy approximations used in process-based vegetation models, first against a high-resolution reference and then through their consequences in a dynamic model. Process-based vegetation models seek to encode core processes in the development of forests, enabling study and prediction of forest successional dynamics, life histories, mechanisms for species coexistence, and carbon-water-nutrient cycles. The latest models simulate the entire life-cycle of individuals, enabling users to address eco-evolutionary questions involving interactions between competing individuals, both from different species or phenotypes within a species (e.g. Weng et al. 2015; Falster et al. 2016; Fisher et al. 2018; Koven et al. 2020). In this context, a model’s ability to accurately capture competition among individuals will strongly influence its predictions.

Comparing the approximations against MAESPA, we found that the widely used PPA failed to capture the response of individual light capture around the point of transition, and that the Flat Top failed in the same way in stands of uniform height: their assumptions impose a stepwise, discontinuous response for individuals varying in height (Fig. 3, Fig. 4), whereas the more complex Deep Crown reproduced the continuous response of the reference. Embedding the same kinds of representation in a dynamic model showed that this discontinuity is not a cosmetic detail. While all approximations developed sensible stands, the PPA’s stepped representation shifted the demographic equilibrium of the population by an order of magnitude and made invasion fitness erratic. Every runnable representation still produced a continuously stable evolutionary endpoint (CSS), so the existence of an endpoint was robust to the approximation; but under the PPA that endpoint was reached through a stepped selection gradient and a fitness curvature about twentyfold steeper, so its location, and the direction of selection between nearby strategies, were not reliably resolved. This holds even though the PPA we tested is a smoothed regularisation of the field’s literal hard-step discretisation, which cannot be integrated by a continuous-time solver at all; the hard-step version would be more severe still. The dynamic experiments further revealed which simplification is responsible: continuity of individual light capture follows from the shape of the shade a plant casts on its neighbours, not from how finely it resolves the light within its own crown. An approximation that casts smooth shade but reads its own light at a single point still yields smooth, usable fitness, whereas one that casts shade as a discrete step does not.

No approximation reproduced every facet of the high-resolution model, and thus none can model a forest system without bias. In particular, all were vulnerable to self-shading – a shared weakness, examined below, that worsens at realistic leaf area – and an approximation could be accurate for aggregate, stand-level light interception while still misrepresenting the variation among individuals that drives competition. Overall, our results reframe the familiar trade-off between detail and computational expense: the property that decides whether an approximation can support competitive and evolutionary inference is the continuity of the shade each plant casts as a function of height. Researchers should match the approximation to the question, and be aware that a representation adequate for one may be badly misleading for another.

### 4.1 Point of transition

The individual competition experiment illustrated how the approximations performed around a point of transition, where the focal tree moves from being slightly taller to slightly shorter than the average tree in the stand. Like the scenarios modelled in the virtual stands, real trees may display small differences in traits or chance growth advantages, leading to scenarios where one tree is only ever slightly taller or shorter than surrounding individuals. When simulated at the point of transition (with zero height variation) both the PPA and the Flat Top approximations failed to capture accurate absorbed PAR, with both grossly overestimating an individual’s light capture if it were slightly taller than the rest of the stand (Fig. 4). The inverse is true when an individual is slightly shorter than the surrounding stand, rendering them highly disadvantaged in the simulation. In a dynamic model, such responses would disproportionately benefit or punish individuals growing in competition with each other. In contrast, the Deep Crown approximation tracked the reference closely, especially in the denser stands (LAI 2.97–5.44 m² m^-2^), where it performed nearly identically to MAESPA; it was less accurate in sparse stands, discussed below.

These different responses emerge directly from the assumptions of each approximation. The two simpler approximations assume no crown overlap between individuals, therefore when an individual is only slightly taller than the stand (or other individuals), its whole leaf area is potentially lifted into higher light (Fig. 1). The simple approximations also assume only one light level for each individual (Moorcroft et al. 2001; Strigul et al. 2008). In stands where heights vary, the Flat Top’s single large step is replaced by many small ones, one at the height of each neighbour, so its response approaches that of Deep Crown; the PPA, which pools neighbours into whole canopy layers, keeps its large steps. In contrast, the Deep Crown assumes that the leaf area of individuals overlaps in the vertical dimension (Falster et al. 2011). Accordingly, when an individual is slightly taller than the surrounding stand, only some of its leaves rise above the other crowns, which is more similar to real systems than the other approximations. Because the Deep Crown slices each crown, each crown can also experience multiple light levels, though at greater computational expense.

Our results reinforce concerns voiced in a review of the previous generation of individual-based “gap” models, where Bugmann (2001) commented that the Flat Top approximation would sharply increase the asymmetry of competition among individuals. Despite the detail devoted to modelling leaf physiological behaviour and radiative schemes in Earth System and vegetation models (see Fisher et al. 2018), many models retain highly simplified representations of competitive shading; these have gone largely untested, and the field has somewhat moved towards simpler representations such as the PPA.

### 4.2 Light beneath open canopies

In sparse stands (LAI 0.44–1.42 m² m^-2^), all of the canopy approximations kept a tree growing up beneath its neighbours in deep shade until it reached their crowns, whereas in MAESPA its light rose earlier and more gradually (Fig. 4). This likely arises because the approximations spread the leaf area of neighbours evenly over the ground, and account for changing hourly PAR without the subtlety of the sun’s angle (Fisher et al. 2018). Beneath widely spaced crowns, a short tree receives direct light through gaps between crowns at some times of day and is shaded at others, depending on the angle of the sun; the dependence of canopy light transmission on the sun’s angle is well documented (Braghiere et al. 2020). Because the light model was fitted to trees at the extremes of shade and full light, this mismatch lies in the shape of the response rather than its end points: in open stands, the approximations exaggerate the disadvantage of trees growing up beneath taller neighbours, and so the steepness of competition among them. Equally, when the canopy is not closed the PPA ignores this competition entirely, placing every tree in full light.

### 4.3 Vulnerability to self-shading

Self-shading occurs as leaves within an individual tree shade each other (Ackerly 1999), an effect that is particularly strong in evergreen trees with extended leaf lifespans and in plants that develop a dense canopy (Ishida et al. 1999; Duursma et al. 2012). Because MAESPA represents the canopy in three dimensions it captures self-shading, whereas none of the approximations do. By deliberately probing this hard case, we found every approximation to be vulnerable: all overestimated light interception at realistic leaf area, and the divergence grew with crown leaf area (Fig. 5). Even Deep Crown, the most accurate approximation for competition, was affected. This vulnerability is therefore a property shared across the approximations rather than one that distinguishes them, and it is orthogonal to the shade-cast continuity that governs competition: self-shading sets the absolute level of light capture, and hence productivity, but not the continuity of the response that fitness requires. It nonetheless matters, and identifies a clear target for future work. Self-shading has so far been added to Deep Crown-type representations only through costly precomputation of crown openness (Lindh et al. 2018) or a simple within-crown treatment (Fischer et al. 2016), and our results reinforce Fisher et al. (2018), who highlighted shading within individuals as a major area of development needed in large-scale vegetation models: any large-scale model using these approximations should incorporate self-shading, or it will overestimate light capture.

### 4.4 Implications for multi-species and trait-based models

Although we examined a single species, the consequences extend directly to the multi-species and trait-based models increasingly used to address eco-evolutionary questions, in which many strategies compete and selection acts on their traits (e.g. Fisher et al. 2018; Koven et al. 2020). For strategies that differ markedly, a discretised representation is largely harmless: the fitness differences between them dwarf the noise introduced at layer boundaries, so the coexistence or exclusion of distinct types is captured well. The difficulty is confined to the opposite and most delicate case – discriminating strategies that differ only slightly. Two near-identical strategies can fall on opposite sides of a layer boundary and receive incomparable fitnesses, so the selection gradient between them – the quantity that determines which way a trait evolves – becomes erratic. A model can therefore produce sensible assemblage-scale simulations and resolve clearly distinct strategies, yet remain unable to say which of two similar strategies wins. This is precisely the regime that matters for predicting trait evolution and fine-scale coexistence. Consistent with this sensitivity, competitive outcomes among coexisting types in this family of models depend strongly on the rules of height-based light competition (Koven et al. 2020), and robustly reproducing multi-type coexistence remains a persistent challenge (Li et al. 2023). None of this means that discrete representations such as the PPA or the Flat Top are wrong for the purposes they were built for. Large models implement them as hard steps – canopy layers or thin crowns – advanced on a fixed timestep, which always returns an answer – but an answer is not necessarily a converged solution. When we forced the hard-step PPA through a fixed-step integrator, a resident’s seed output did not converge as the timestep was refined, varying about tenfold with the step size, whereas the smoothed PPA converged (Fig. **S8**); the hard-step Flat Top casts an equally discontinuous shade. The difficulty is therefore present in fixed-step models too, though hidden; and it bears most heavily on resolving selection among similar strategies, where continuity of the cast shade – absent from a hard-layered profile – is what the inference requires.

### 4.5 Limitations

Our two-stage design lets each model do what the other cannot: MAESPA supplies a high-resolution benchmark for instantaneous light capture, and the dynamic model carries those differences forward to lifetime and evolutionary outcomes. The two stages are nonetheless run in different models, and because MAESPA is static we could not validate the dynamic predictions directly against a high-resolution model that simulates growth; closing this gap would require a high-resolution model capable of simulating stand development through time. Second, we examined a single species and a single trait axis. This is sufficient to expose the behaviour of each approximation, but leaves the multi-species case – where many strategies compete and selection acts on their traits – to be addressed by extension, as discussed above. Third, the virtual stands are stylised: trees sit on a regular grid, crowns of identical size overlap in the densest stands, and all three approximations share one empirical light model fitted to MAESPA. These choices hold everything but the canopy representation fixed, but the absolute errors we report depend on them. Fourth, we considered only light interception. Beyond light interception, mechanistic models calculate carbon uptake, photosynthesis, carbon budgets, transpiration and more; extending the comparison from absorbed PAR to these outputs would further test how the approximations diverge from high-resolution models and real forest systems.

### 4.6 Closing remarks

Our study highlights the trade-offs inherent in approximating canopies, and refines where the important trade-off actually lies. Greater complexity can buy greater fidelity: the more detailed Deep Crown best captured individual competition, and embedding it in a dynamic model gave smooth, well-behaved fitness. But our results show that the decisive property is not overall complexity or accuracy – it is the continuity of the shade each plant casts as a function of height. An approximation that preserves that continuity can be coarse elsewhere, including in how it resolves a plant’s own light capture, and still support competitive and evolutionary inference; one that breaks it cannot, however else it is refined. The cost differences are real: both the PPA and the Flat Top require only one calculation of light capture per crown per timestep, whereas our implementation of Deep Crown requires 50, and MAESPA typically calculates radiation at 72 grid points in each target crown, each resolved into sunlit and shaded leaf area (Duursma and Medlyn 2012). That the approximations approach MAESPA at a small fraction of this cost explains their central role in current models, where high-resolution models would demand more computation than is feasible. Nonetheless, our results expose limitations in the approximations now used in large-scale models – most pressingly their discontinuous treatment of competitive shading and their neglect of self-shading – and motivate new approximations that preserve continuity of the cast shade, and represent self-shading, without much additional expense.

## Acknowledgements

This work was supported by an Australian Research Council Future Fellowship to D. Falster (FT160100113). D. Falster is supported by ARC grant (LP230100049). We thank B. Medlyn and M. Sabot for helpful suggestions and I. Towers for comments on an earlier draft.

## CRediT authorship contribution statement

**Lily P Dun**: Conceptualisation, Data curation, Formal analysis, Investigation, Methodology, Software, Visualisation, Writing – original draft. **Daniel S Falster**: Conceptualisation, Formal analysis, Funding acquisition, Methodology, Writing – review and editing.

## Data Availability

All data generated in the MAESPA simulations, along with code used to generate stands and analyse results, are archived on Zenodo (Dun and Falster, 2026; https://doi.org/10.5281/zenodo.24334797). The dynamic-model experiments used the plant package (https://github.com/traitecoevo/plant); the code and cached outputs for these experiments are in that repository at commit 81cfeab (vignettes/examples/DunFalster2026/).

## Declaration of generative AI use

During the preparation of this work, the authors used Claude (Anthropic) for coding support in the analysis and figure scripts. After using this tool, the authors reviewed and edited the content as needed and take full responsibility for the content of the published article.

## References

Ackerly, D. 1999. “Self-Shading, Carbon Gain and Leaf Dynamics: A Test of Alternative Optimality Models.” Oecologia 119 (3): 300–310. 10.1007/s004420050790.

Beaudet, M., C. Messier, and C. D. Canham. 2002. “Predictions of Understory Light Conditions in Northern Hardwood Forests Following Parameterization, Sensitivity Analysis, and Tests of the SORTIE Light Model.” Forest Ecology and Management 165: 235–48. 10.1016/S0378-1127(01)00621-1.

Braghiere, Renato K., Tristan Quaife, Emily Black, et al. 2020. “Influence of Sun Zenith Angle on Canopy Clumping and the Resulting Impacts on Photosynthesis.” Agricultural and Forest Meteorology 291: 108065. 10.1016/j.agrformet.2020.108065.

Braghiere, Renato K., Tristan Quaife, Emily Black, Liang He, and Jing M. Chen. 2019. “Underestimation of Global Photosynthesis in Earth System Models Due to Representation of Vegetation Structure.” Global Biogeochemical Cycles 33 (11): 1358–69. 10.1029/2018GB006135.

Bugmann, Harald. 2001. “A Review of Forest Gap Models.” Climatic Change 51 (3–4): 259–305. 10.1023/A:1012525626267.

Christina, M., Y. Nouvellon, J. P. Laclau, et al. 2017. “Importance of Deep-Water Uptake in Tropical Eucalypt Forest.” Functional Ecology 31: 509–19. 10.1111/1365-2435.12727.

Christina, M., Y. Nouvellon, J. P. Laclau, O. C. Campoe, and G. le Maire. 2016. “Sensitivity and Uncertainty Analysis of the Carbon and Water Fluxes at the Tree Scale in Eucalyptus Plantations Using a Metamodeling Approach.” Canadian Journal of Forest Research 46: 297–309. 10.1139/cjfr-2015-0173.

Cox, P. M., R. A. Betts, C. B. Bunton, R. L. H. Essery, P. R. Rowntree, and J. Smith. 1999. “The Impact of New Land Surface Physics on the GCM Simulation of Climate and Climate Sensitivity.” Climate Dynamics 15: 183–203. 10.1007/s003820050276.

Duursma, R. A. 2015. Maeswrap: Wrapper Functions for MAESTRA/MAESPA. 10.32614/CRAN.package.Maeswrap.

Duursma, R. A., Daniel S. Falster, F. Valladares, et al. 2012. “Light Interception Efficiency Explained by Two Simple Variables: A Test Using a Diversity of Small-to Medium-Sized Woody Plants.” New Phytologist 193 (2): 397–408. 10.1111/j.1469-8137.2011.03943.x.

Duursma, R. A., and A. Mäkelä. 2007. “Summary Models for Light Interception and Light-Use Efficiency of Non-Homogeneous Canopies.” Tree Physiology 27 (6): 859–70. 10.1093/treephys/27.6.859.

Duursma, R. A., and B. E. Medlyn. 2012. “MAESPA: A Model to Study Interactions Between Water Limitation, Environmental Drivers and Vegetation Function at Tree and Stand Levels, with an Example Application to [CO_2_] × Drought Interactions.” Geoscientific Model Development 5 (4): 919–40. 10.5194/gmd-5-919-2012.

Eckes-Shephard, Annemarie H., Arthur P. K. Argles, Bogdan Brzeziecki, et al. 2025. “Demography, Dynamics and Data: Building Confidence for Simulating Changes in the World’s Forests.” New Phytologist 248 (6): 2722–49. 10.1111/nph.70643.

Falster, Daniel S., Å. Brännström, U. Dieckmann, and M. Westoby. 2011. “Influence of Four Major Plant Traits on Average Height, Leaf-Area Cover, Net Primary Productivity, and Biomass Density in Single-Species Forests: A Theoretical Investigation.” Journal of Ecology 99 (1): 148–64. 10.1111/j.1365-2745.2010.01735.x.

Falster, Daniel S., Åke Brännström, Mark Westoby, and Ulf Dieckmann. 2017. “Multitrait Successional Forest Dynamics Enable Diverse Competitive Coexistence.” Proceedings of the National Academy of Sciences 114 (13): E2719–28. 10.1073/pnas.1610206114.

Falster, Daniel S., Remko A. Duursma, Masae I. Ishihara, et al. 2015. “BAAD: A Biomass and Allometry Database for Woody Plants.” Ecology 96 (5): 1445. 10.1890/14-1889.1.

Falster, Daniel S., Richard G. FitzJohn, Åke Brännström, Ulf Dieckmann, and Mark Westoby. 2016. “Plant: A Package for Modelling Forest Trait Ecology and Evolution.” Methods in Ecology and Evolution 7 (2): 136–46. 10.1111/2041-210X.12525.

Falster, Daniel S., and M. Westoby. 2003. “Plant Height and Evolutionary Games.” Trends in Ecology and Evolution 18 (7): 337–43. 10.1016/S0169-5347(03)00061-2.

Fischer, Rico, Friedrich Bohn, Mateus Dantas de Paula, et al. 2016. “Lessons Learned from Applying a Forest Gap Model to Understand Ecosystem and Carbon Dynamics of Complex Tropical Forests.” Ecological Modelling 326: 124–33. 10.1016/j.ecolmodel.2015.11.018.

Fisher, R. A., S. Muszala, M. Verteinstein, et al. 2015. “Taking Off the Training Wheels: The Properties of a Dynamic Vegetation Model Without Climate Envelopes, CLM4.5(ED).” Geoscientific Model Development 8 (11): 3593–619. 10.5194/gmd-8-3593-2015.

Fisher, Rosie A., Charles D. Koven, William R. L. Anderegg, et al. 2018. “Vegetation Demographics in Earth System Models: A Review of Progress and Priorities.” Global Change Biology 24 (1): 35–54. 10.1111/gcb.13910.

Fisher, Rosie, Nate McDowell, Drew Purves, et al. 2010. “Assessing Uncertainties in a Second-Generation Dynamic Vegetation Model Caused by Ecological Scale Limitations.” New Phytologist 187 (3): 666–81. 10.1111/j.1469-8137.2010.03340.x.

Geritz, S. A. H., É. Kisdi, G. Meszéna, and J. A. J. Metz. 1998. “Evolutionarily Singular Strategies and the Adaptive Growth and Branching of the Evolutionary Tree.” Evolutionary Ecology 12 (1): 35–57. 10.1023/A:1006554906681.

Huston, M., and T. Smith. 1987. “Plant Succession: Life History and Competition.” The American Naturalist 130 (2): 168–98. 10.1086/284704.

Ishida, A., A. Uemura, N. Koike, Y. Matsumoto, and A. L. Hoe. 1999. “Interactive Effects of Leaf Age and Self-Shading on Leaf Structure, Photosynthetic Capacity and Chlorophyll Fluorescence in the Rain Forest Tree, *Dryobalanops Aromatica*.” Tree Physiology 19: 741–47. 10.1093/treephys/19.11.741.

Kim, H.-S., S. Palmroth, M. Therezien, P. Stenberg, and R. Oren. 2011. “Analysis of the Sensitivity of Absorbed Light and Incident Light Profile to Various Canopy Architecture and Stand Conditions.” Tree Physiology 31 (1): 30–47. 10.1093/treephys/tpq098.

Koven, Charles D., Ryan G. Knox, Rosie A. Fisher, et al. 2020. “Benchmarking and Parameter Sensitivity of Physiological and Vegetation Dynamics Using the Functionally Assembled Terrestrial Ecosystem Simulator (FATES) at Barro Colorado Island, Panama.” Biogeosciences 17 (11): 3017–44. 10.5194/bg-17-3017-2020.

Kükenbrink, D., F. D. Schneider, B. Schmid, J. P. Gastellu-Etchegorry, M. E. Schaepman, and F. Morsdorf. 2021. “Modelling of Three-Dimensional, Diurnal Light Extinction in Two Contrasting Forests.” Agricultural and Forest Meteorology 296: 108230. 10.1016/j.agrformet.2020.108230.

Li, Lingcheng, Yilin Fang, Zhonghua Zheng, et al. 2023. “A Machine Learning Approach Targeting Parameter Estimation for Plant Functional Type Coexistence Modeling Using ELM-FATES (V2.0).” Geoscientific Model Development 16 (14): 4017–40. 10.5194/gmd-16-4017-2023.

Lindh, Magnus, Daniel S. Falster, Lai Zhang, Ulf Dieckmann, and Åke Brännström. 2018. “Latitudinal Effects on Crown Shape Evolution.” Ecology and Evolution 8 (16): 8149–58. 10.1002/ece3.4275.

Luo, Yiqi, Belinda Medlyn, Dafeng Hui, David Ellsworth, James Reynolds, and Gabriel Katul. 2001. “Gross Primary Productivity in Duke Forest: Modeling Synthesis of CO_2_ Experiment and EddyFlux Data.” Ecological Applications 11 (1): 239–52. https://doi.org/10.1890/1051-0761(2001)011%5B0239:GPPIDF%5D2.0.CO;2.

Maxwell, J. D., A. C. Rhodes, and S. B. St. Clair. 2019. “Human Altered Disturbance Patterns and Forest Succession: Impacts of Competition and Ungulate Herbivory.” Oecologia 189: 1061–70. 10.1007/s00442-019-04370-8.

Medlyn, B. E. 2004. “A MAESTRO Retrospective.” In Forests at the Land-Atmosphere Interface, edited by M. Mencuccini, J. Grace, J. B. Moncrieff, and K. G. McNaughton. CAB International. 10.1079/9780851996776.0105.

Metz, J. A. J., R. M. Nisbet, and S. A. H. Geritz. 1992. “How Should We Define ‘Fitness’ for General Ecological Scenarios?” Trends in Ecology and Evolution 7 (6): 198–202. 10.1016/0169-5347(92)90073-K.

Moorcroft, P. R., G. C. Hurtt, and S. W. Pacala. 2001. “A Method for Scaling Vegetation Dynamics: The Ecosystem Demography Model (ED).” Ecological Monographs 71 (4): 557–86. https://doi.org/10.1890/0012-9615(2001)071%5B0557:AMFSVD%5D2.0.CO;2.

Norman, J. M. 1979. “Modeling the Complete Crop Canopy.” In Modification of the Aerial Environment of Plants, edited by B. J. Barfield and J. F. Gerber. ASAE Monograph 2. American Society of Agricultural Engineers.

Norman, J. M. 1993. “Scaling Processes Between Leaf and Canopy Levels.” In Scaling Physiological Processes: Leaf to Globe, edited by J. R. Ehleringer and C. B. Field. Academic Press. 10.1016/B978-0-12-233440-5.50010-5.

Pacala, S. W., and D. H. Deutschman. 1995. “Details That Matter: The Spatial Distribution of Individual Trees Maintains Forest Ecosystem Function.” Oikos 74 (3): 357–65. 10.2307/3545980.

Pearcy, R. W., and W. Yang. 1996. “A Three-Dimensional Crown Architecture Model for Assessment of Light Capture and Carbon Gain by Understory Plants.” Oecologia 108 (1): 1–12. 10.1007/BF00333208.

Purves, Drew, and Stephen Pacala. 2008. “Predictive Models of Forest Dynamics.” Science 320 (5882): 1452–53. 10.1126/science.1155359.

R Core Team. 2026. R: A Language and Environment for Statistical Computing. R Foundation for Statistical Computing. https://www.R-project.org/.

Reis, Mariana Gonçalves dos, and Aristides Ribeiro. 2020. “Conversion Factors and General Equations Applied in Agricultural and Forest Meteorology.” Agrometeoros 27 (2): 227–58. 10.31062/agrom.v27i2.26527.

Rogers, Alistair, Belinda E. Medlyn, Jeffrey S. Dukes, et al. 2017. “A Roadmap for Improving the Representation of Photosynthesis in Earth System Models.” New Phytologist 213 (1): 22–42. 10.1111/nph.14283.

Roupsard, O., J. Dauzat, Y. Nouvellon, et al. 2008. “Cross-Validating Sunshade and 3D Models of Light Absorption by a Tree-Crop Canopy.” Agricultural and Forest Meteorology 148: 549–64.10.1016/j.agrformet.2007.11.002.

Salas-Eljatib, Christian, and Aaron R. Weiskittel. 2018. “Evaluation of Modeling Strategies for Assessing Self-Thinning Behavior and Carrying Capacity.” Ecology and Evolution 8 (22): 10768–79. 10.1002/ece3.4525.

Song, C., and L. E. Band. 2004. “MVP: A Model to Simulate the Spatial Patterns of Photosynthetically Active Radiation Under Discrete Forest Canopies.” Canadian Journal of Forest Research 34: 1192–203. 10.1139/x03-280.

Stadt, K. J., and V. G. Lieffers. 2000. “MIXLIGHT: A Flexible Light Transmission Model for Mixed-Species Forest Stands.” Agricultural and Forest Meteorology 102: 235–52. 10.1016/S0168-1923(00)00128-3.

Strigul, Nikolay, Denis Pristinski, Drew Purves, Jonathan Dushoff, and Stephen Pacala. 2008. “Scaling from Trees to Forests: Tractable Macroscopic Equations for Forest Dynamics.” Ecological Monographs 78 (4): 523–45. 10.1890/08-0082.1.

Vezy, R., M. Christina, O. Roupsard, et al. 2018. “Measuring and Modelling Energy Partitioning in Canopies of Varying Complexity Using MAESPA Model.” Agricultural and Forest Meteorology 253–254: 203–17. 10.1016/j.agrformet.2018.02.005.

Wang, Y. P., and P. G. Jarvis. 1990. “Influence of Crown Structural Properties on PAR Absorption, Photosynthesis, and Transpiration in Sitka Spruce: Application of a Model (MAESTRO).” Tree Physiology 7 (1-4): 297–316. 10.1093/treephys/7.1-2-3-4.297.

Wang, Y. P., A. Rey, and P. G. Jarvis. 1998. “Carbon Balance of Young Birch Trees Grown in Ambient and Elevated Atmospheric CO_2_ Concentrations.” Global Change Biology 4 (8): 797–807. 10.1046/j.1365-2486.1998.00170.x.

Weng, E. S., S. Malyshev, J. W. Lichstein, et al. 2015. “Scaling from Individual Trees to Forests in an Earth System Modeling Framework Using a Mathematically Tractable Model of Height-Structured Competition.” Biogeosciences 12 (9): 2655–94. 10.5194/bg-12-2655-2015.

Westoby, M. 1984. “The Self-Thinning Rule.” Advances in Ecological Research 14: 167–225. 10.1016/S0065-2504(08)60171-3.

Woodgate, W. 2013. Tumbarumba OzFlux Tower Site. OzFlux: Australian and New Zealand Flux Research and Monitoring. Online data collection. https://data.ozflux.org.au/portal/pub/viewColDetails.jspx?collection.id=1882717.

Yokozawa, M., and T. Hara. 1995. “Foliage Profile, Size Structure and Stem Diameter–Plant Height Relationship in Crowded Plant Populations.” Annals of Botany 76 (3): 271–85. 10.1006/anbo.1995.1096.

