## Supplemental material for "Continuity, not accuracy: how canopy approximations shape competition and selection in size-structured vegetation models"

Lily P Dun Daniel S Falster

This document contains supporting figures and tables referenced in the main text. **Figures S1–S5** and **Table S1** describe the MAESPA computational experiments (Stage 1); **Tables S2–S3** and **Figures S6–S8** describe the dynamic-model experiments (Stage 2, `plant`).

#### Stage 1 — MAESPA computational experiments

#### Table S1. Trees and stands in the MAESPA experiments

All trees other than the focal tree and the edge-effect wall had the same crown and leaf area; only trunk height varied, drawn from a normal distribution with mean 15 m and the stated coefficient of variation (CV).

| Parameter | Value |
|---|---|
| Crown radius, x and y directions (m) | 2.54 |
| Crown height (depth) (m) | 6.36 |
| Mean trunk height (m) | 15 |
| Leaf area per tree ($m^2$) | 27 |
| Plot size (m) | 196 × 196 |
| Edge-effect wall | two outermost rows on each side; leaf area 2,000 $m^2$, crown radius 13 m, crown from near ground to 21.4 m |

| Trees per row | Tree spacing (m) | Stand LAI ($m^2$ $m^{-2}$) | Crown cover (crown area per ground area) |
|---|---|---|---|
| 26 | 7.84 | 0.44 | 0.33 |
| 46 | 4.36 | 1.42 | 1.07 |
| 66 | 3.02 | 2.97 | 2.24 |
| 79 | 2.51 | 4.28 | 3.22 |
| 89 | 2.23 | 5.44 | 4.10 |

**Figure S1.** Design and workflow of the two computational experiments: (**a**) individual competition, varying the stand's height variation, the focal tree's height relative to the stand and the stand's LAI; (**b**) vulnerability to self-shading, varying the focal tree's height and leaf area in a single stand. Each stand was simulated in MAESPA, and the leaf area above the focal tree under each canopy approximation was converted to absorbed PAR with a light model fitted to MAESPA (see Methods).

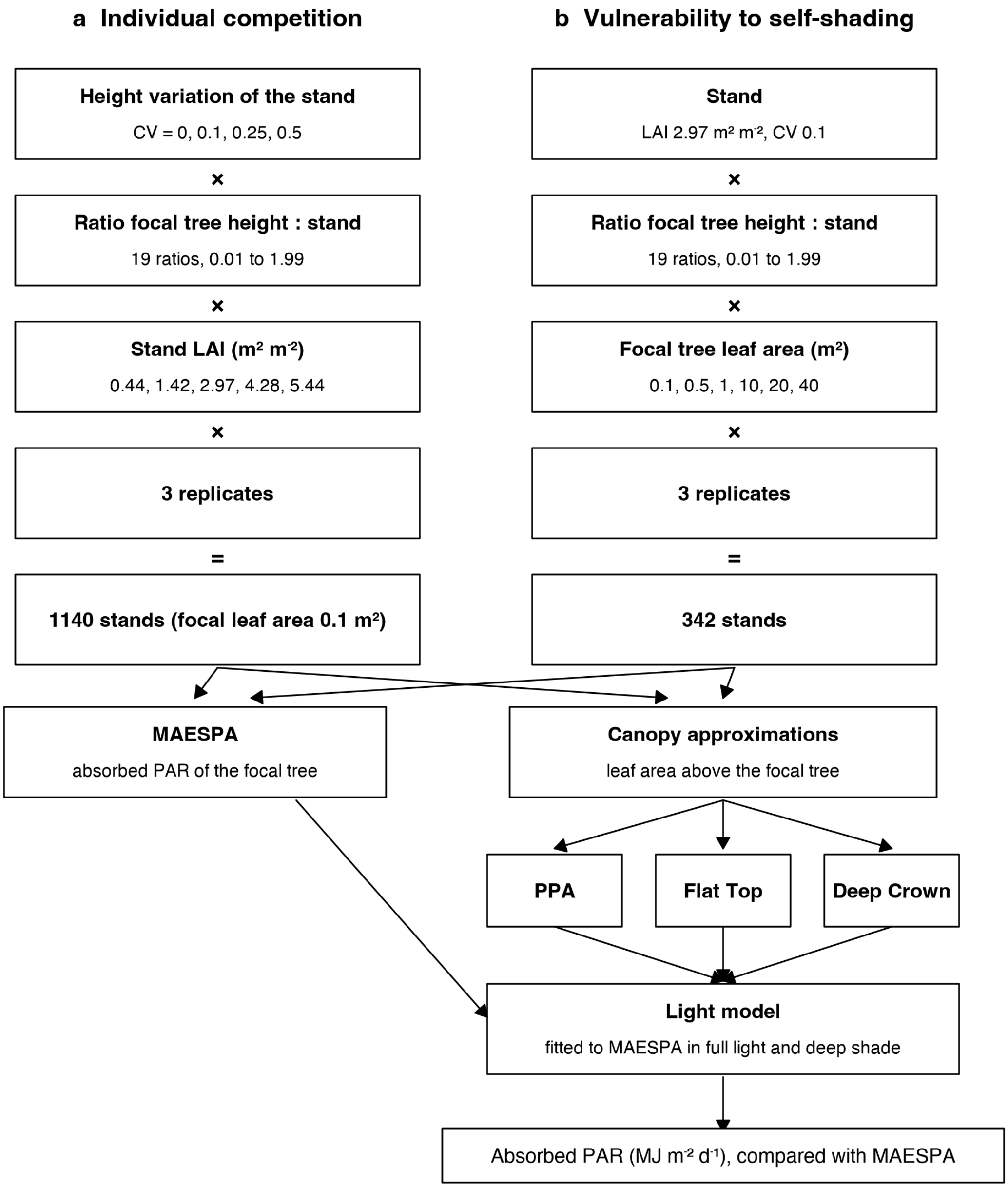

**Figure S2.** A whole MAESPA stand (LAI 1.42 $m^2$ $m^{-2}$, no height variation, focal tree in red), including the wall of large trees around the plot edge used to reduce edge effects near sunrise and sunset.

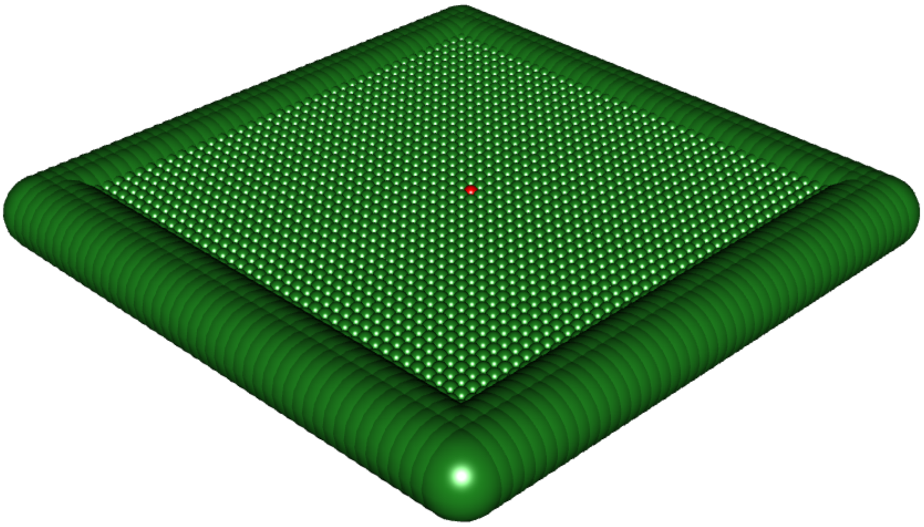

**Figure S3.** The simulated tree (red) against the relationships between total height and leaf area, trunk height and crown width in the BAAD database (Falster et al. 2015). Each point is an individual plant; green points are *Eucalyptus*. Axes are log-scaled. The tree's leaf area and crown depth are typical of *Eucalyptus* of similar height; its crown is wider than typical.

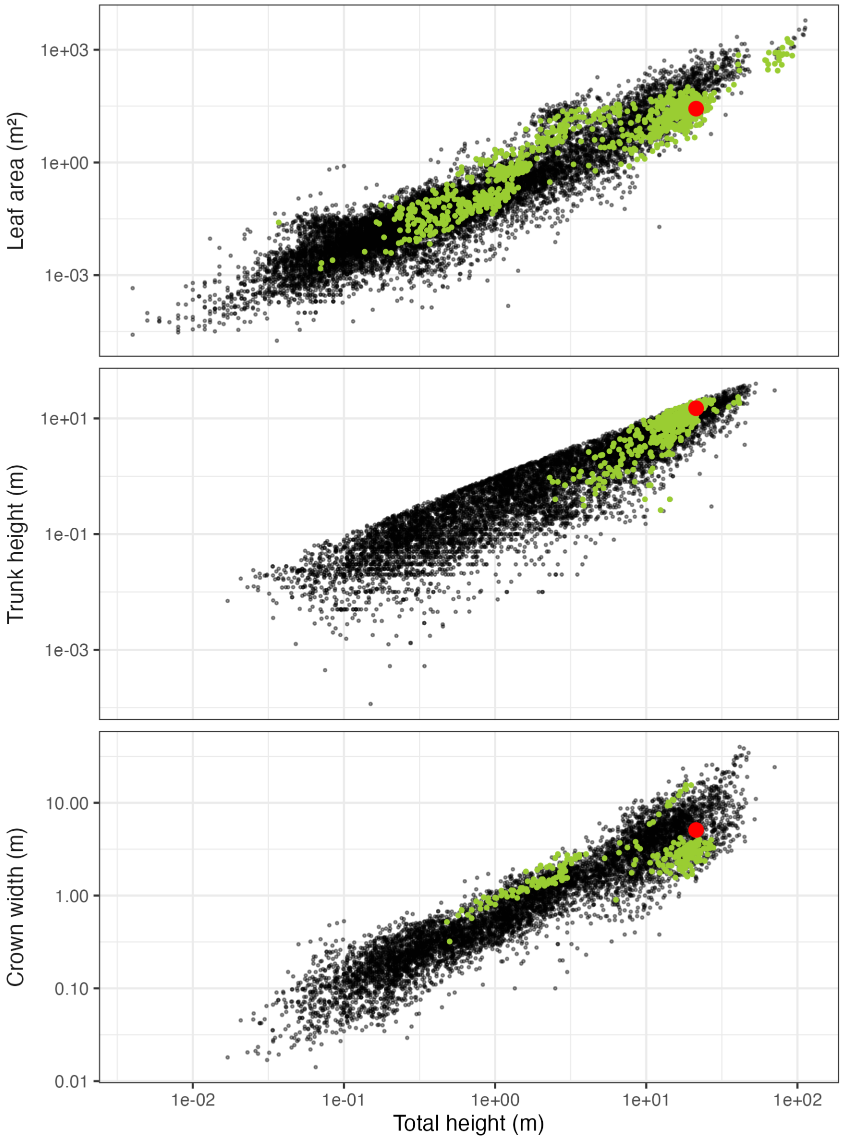

**Figure S4.** Comparison of light models. Absorbed PAR for a focal tree against its height ratio to the stand, for MAESPA and for Deep Crown with a single-component Beer's law (dashed) and with the two-component light model used in the paper (solid), each fitted to MAESPA in the same way (see Methods). Stand LAI 2.97 $m^2$ $m^{-2}$, CV 0.1. A single exponential cannot match MAESPA's deep shade across stand densities: fitted across all densities, it makes shaded trees too dark here, and far too dark in the densest stands (0.04 against 0.26 MJ $m^{-2}$ $d^{-1}$ in MAESPA at LAI 5.44).

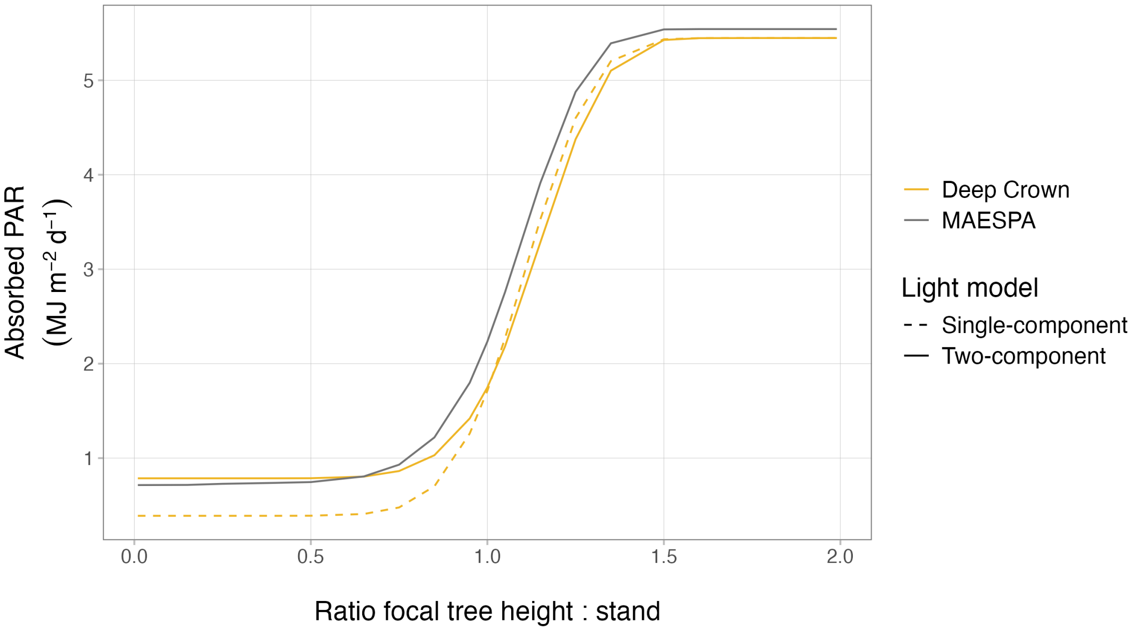

**Figure S5.** Vulnerability to self-shading across the full range of focal leaf areas (0.1–40 $m^2$). Absorbed PAR for a focal tree against its height ratio to the stand, for MAESPA and the three approximations, at each focal leaf area; the approximations increasingly overestimate light capture as focal leaf area grows, because none represent self-shading. Stand LAI 2.97 $m^2$ $m^{-2}$ and CV 0.1. MAESPA lines join the 19 simulated height ratios; the approximations are drawn at height ratios 0.005 apart. (The main-text self-shading figure shows the 0.1 and 40 $m^2$ cases.)

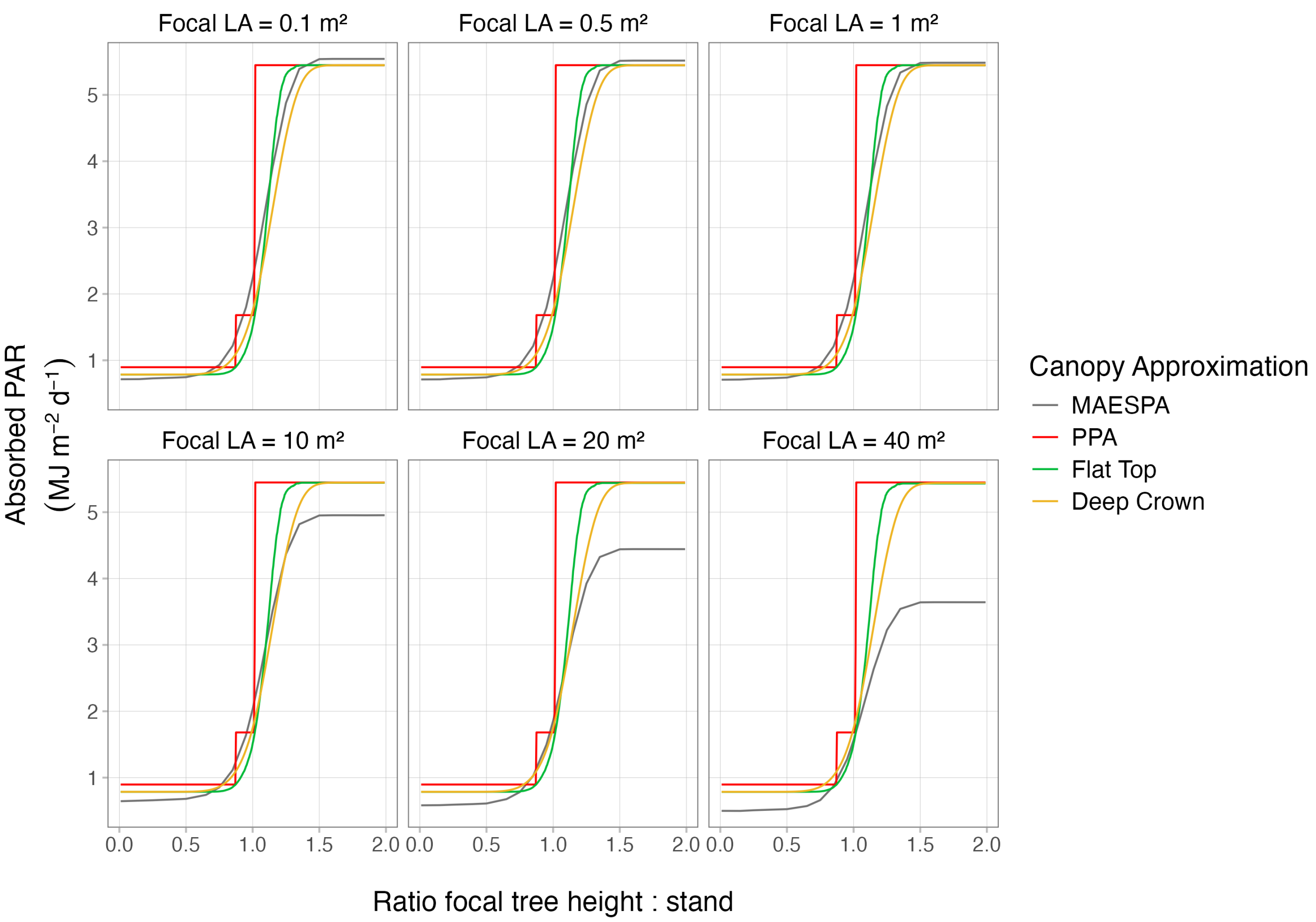

### Stage 2 — dynamic-model experiments

#### Table S2. Canopy representations: paper names and `plant` settings

The names used in the main text, what each represents along the two axes (the shade a plant casts, and how it reads its own light), and the corresponding `plant shading_model` setting.

| Paper name | Meaning | `plant` setting |
|---|---|---|
| Deep Crown | smooth (Yokozawa) shade; assimilation integrated over crown depth | `deep-crown` |
| Mean-Light | smooth shade; light averaged over crown depth, one assimilation evaluation | `mean-light` |
| Crown-Centre | smooth shade; one assimilation evaluation at the crown centre | `crown-centre` |
| PPA (smoothed) | layered, stepped light profile, boundaries smoothed; runnable, used for all PPA results | `ppa`, `ppa_layer_smoothing` > 0 (default 0.3) |
| PPA (hard step) | literal field discretisation; discontinuous light profile, does not run in the adaptive solver | `ppa`, `ppa_layer_smoothing` = 0 |
| Flat Top (smoothed) | box / thin-layer shade, smoothed into a continuous drop; runs but biased | `flat-top-soft-box` |
| Flat Top (hard step) | box / thin-layer shade as a hard step; discontinuous, does not run | `flat-top-box` |

The smooth-shade / crown-centre-assimilation model (`crown-centre`) is **not** the field's "Flat Top": "Flat Top" denotes the box / thin-layer *shade*. Crown-Centre and Flat Top share the same crown-centre assimilation and differ only in the shade they cast, so contrasting them isolates the shade cast from the assimilation rule.

#### Table S3. Classification of singular strategies

For each representation we located the singular strategy (the trait value at which the selection gradient vanishes) and classified it by convergence stability (sign of the selection-gradient slope) and evolutionary stability (sign of the fitness curvature). All five are continuously stable strategies (CSS): convergence-stable, uninvadable fitness maxima, i.e. genuine evolutionary endpoints rather than branching points. The PPA's CSS is reached through a stepped selection gradient with far steeper curvature and is therefore not robustly located.

| Representation | Singular LMA (kg $m^{-2}$) | Class | Fitness curvature | Robust |
|---|---|---|---|---|
| Deep Crown | 0.1007 | CSS | $-1.2 \times 10^4$ | yes |
| Mean-Light | 0.0976 | CSS | $-1.7 \times 10^4$ | yes |
| Crown-Centre | 0.0924 | CSS | $-2.1 \times 10^4$ | yes |
| Flat Top (smoothed) | 0.0901 | CSS | $-3.1 \times 10^4$ | yes |
| PPA (smoothed) | 0.0859 | CSS | $-2.5 \times 10^5$ | no (stepped gradient) |

**Figure S6.** Size-structure development of a patch under four representations: Deep Crown, Mean-Light, Crown-Centre and PPA (smoothed). Each line is a cohort's height trajectory, with opacity scaled by cohort density; the PPA's discrete layering produces visibly different trajectories.

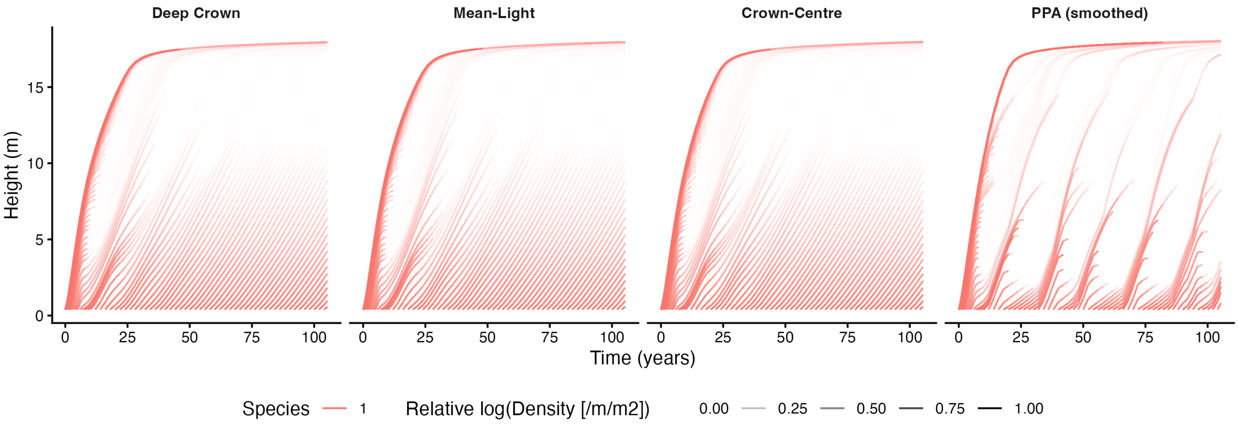

**Figure S7.** Vertical light profile (cumulative leaf area above each height) at several patch ages under four representations: Deep Crown, Mean-Light, Crown-Centre and PPA (smoothed). The three continuous models share a smooth profile; the PPA shows the characteristic staircase, constant within a layer and jumping at each boundary.

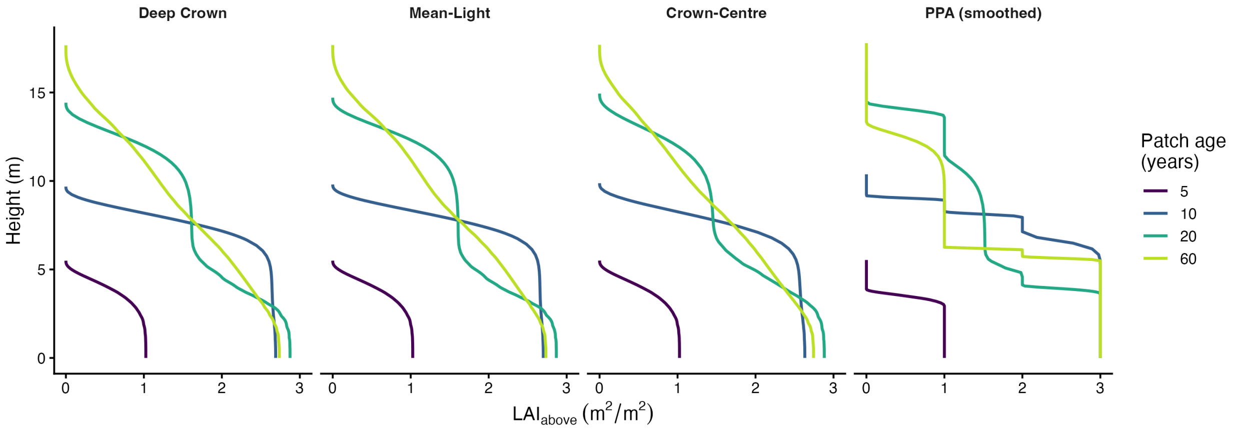

**Figure S8.** The hard-step PPA does not converge when forced through a fixed-step integrator. The adaptive solver used in the main text cannot integrate the hard-step PPA (`ppa_layer_smoothing` $= 0$), whose light profile is discontinuous. Here it is forced to run with fixed-step forward Euler integration, as in large vegetation models. (**a**) Selection gradient against resident LMA for the smoothed PPA (daily steps) and the hard-step PPA (daily and half-daily steps); the smoothed PPA has a single zero crossing, whereas the hard-step gradient is erratic and its sign pattern changes when the step is halved (axis clipped; $|\text{gradient}|$ reaches about 9000). (**b**) Seed output of a single resident (LMA 0.1) as the step is refined: the smoothed PPA converges, whereas the hard-step output varies about tenfold with step size and does not converge (note the different y-axis scales).

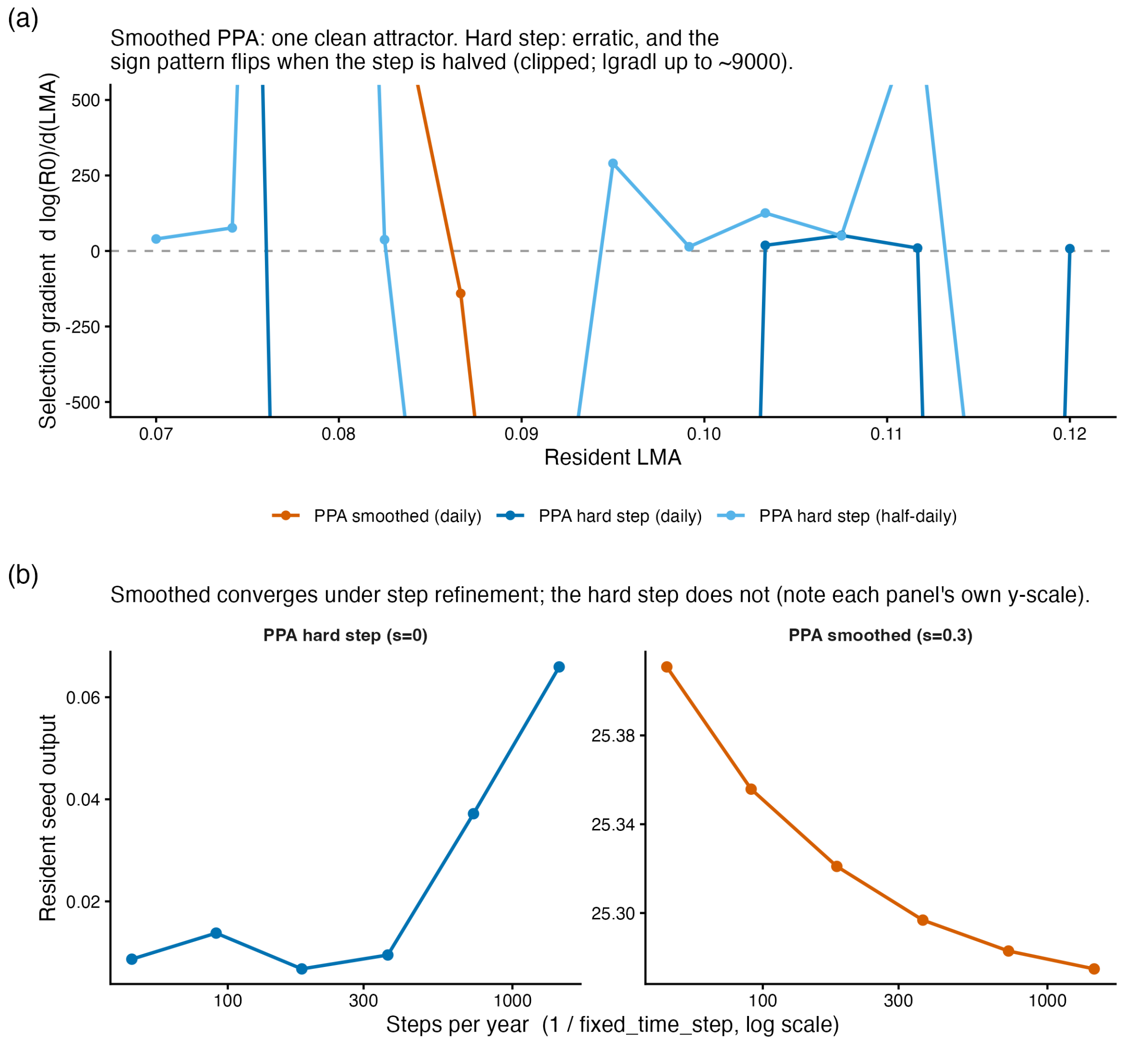

### References

Falster, Daniel S., Remko A. Duursma, Masae I. Ishihara, et al. 2015. “BAAD: A Biomass and Allometry Database for Woody Plants.” *Ecology* 96 (5): 1445. https://doi.org/10.1890/14-1889.1.
